# Integrative genomics sheds light on the immunobiology of tuberculosis in cattle

**DOI:** 10.1101/2024.02.27.582295

**Authors:** John F. O’Grady, Gillian P. McHugo, James A. Ward, Thomas J. Hall, Sarah L. Faherty O’Donnell, Carolina N. Correia, John A. Browne, Michael McDonald, Eamonn Gormley, Valentina Riggio, James G. D. Prendergast, Emily L. Clark, Hubert Pausch, Kieran G. Meade, Isobel C. Gormley, Stephen V. Gordon, David E. MacHugh

## Abstract

*Mycobacterium bovis* causes bovine tuberculosis (bTB), an infectious disease of cattle that poses a zoonotic threat to humans. Research has shown that bTB susceptibility is a heritable trait, and that the peripheral blood (PB) transcriptome is perturbed during bTB disease. Hitherto, no study has integrated PB transcriptomic, genomic and GWAS data to study bTB disease, and little is known about the genomic architecture underpinning the PB transcriptional response to *M. bovis* infection. Here, we perform transcriptome profiling of PB from 63 control and 60 confirmed *M. bovis* infected animals and detect 2,592 differently expressed genes that perturb multiple immune response pathways. Leveraging imputed genome-wide SNP data, we characterise thousands of *cis*- and *trans*-expression quantitative trait loci (eQTLs) and show that the PB transcriptome is substantially impacted by intrapopulation genomic variation. We integrate our gene expression data with summary statistics from multiple GWAS data sets for bTB susceptibility and perform the first transcriptome-wide association study (TWAS) in the context of tuberculosis disease. From this TWAS, we identify 136 functionally relevant genes (including *RGS10*, *GBP4*, *TREML2*, and *RELT*) and provide important new omics data for understanding the host response to mycobacterial infections that cause tuberculosis in mammals.

## Introduction

Tuberculosis (TB) is a chronic infectious disease and a major source of ill health globally with over one billion people having died as a consequence of human TB (hTB) during the past two centuries^1^ and with a further 1.3 million deaths reported in 2022^2^, illustrating both the historical and persistent threat of the disease. The primary causative agent of hTB, *Mycobacterium tuberculosis*, forms part of the *Mycobacterium tuberculosis* complex (MTBC), a group of phylogenetically closely related bacteria exhibiting extreme genomic homogeneity that cause TB disease in mammals^3–6^. Another member of the MTBC, *Mycobacterium bovis*, is the chief causative agent of bovine tuberculosis (bTB), an endemic disease principally associated with cattle that imposes a significant economic impact on individual farmers and national economies^7,8^. As a zoonotic pathogen, *M. bovis* can transmit from animals to humans causing zoonotic TB (zTB), which disproportionally affects the Global South^9,10^. The most recent estimates, available for 2019, attributed more than 140,000 of new hTB cases and more than 11,000 deaths to zTB^11^.

Previous research has shown that there are many shared characteristics between the pathogenesis of hTB and bTB, such that cattle can serve as a valuable large animal model to study TB disease in humans^12–15^. The primary route of infection for both *M. tuberculosis* and *M. bovis* is via the inhalation of aerosolised bacilli expelled by an infected individual or animal that are then phagocytosed by host alveolar macrophages (AM), establishing the primary site of infection in the lung. Normally, efficient pathogen killing is achieved by AMs through a range of innate immune response mechanisms including encasement of the bacilli within a phagolysosome, autophagy and apoptosis of infected cells, and by the production of antimicrobial peptides^16,17^. However, mycobacteria have evolved a range of strategies to manipulate innate immune responses, thereby facilitating colonisation, persistence, and replication within AMs^18–20^. Given the marked genomic similarities between *M. tuberculosis* and *M. bovis*, the close parallels between host-pathogen interactions and disease progression for hTB and bTB, and the zoonotic threat of *M. bovis*, a One Health approach to understanding the molecular mechanisms that underpin host immune responses and pathology in bTB can also provide important new information for tackling both hTB and zTB.

The genetic basis of susceptibility to *M. bovis* infection and bTB disease traits has been examined in cattle using focused candidate gene approaches^21–24^. Previous work has also highlighted the existence of substantial genetic variation for susceptibility to *M. bovis* infection in cattle populations^25,26^. In addition, genome-wide association studies (GWAS) have suggested susceptibility to *M. bovis* infection and bTB disease resilience traits are highly polygenic and influenced by interbreed genetic variation, which is reflected in modest replication of GWAS signals across multiple experiments^27–33^. Ultimately, identifying, cataloguing, and measuring the functional effects of these polymorphisms will expand and enhance genomic prediction models for economically important traits such as resistance to *M. bovis* infection^34^.

Expression quantitative trait loci (eQTLs) are genomic sequence variations—primarily single-nucleotide polymorphisms (SNPs)—that modulate gene expression and mRNA transcript abundance^35–39^. In this regard, SNPs that are significantly associated with a trait of interest often exert an eQTL regulatory effect^40–42^. This is observed for hTB, where infection response eQTLs detected in dendritic cells challenged with *M. tuberculosis* were enriched for SNPs associated with susceptibility to hTB^43^. In cattle, eQTLs and other regulatory polymorphisms have been shown to contribute a substantial proportion of the genetic variation associated with multiple complex traits^44,45^. A transcriptome wide association study (TWAS) is a multi-omics integrative strategy that combines gene expression data and independently generated GWAS summary data to discern explanatory links between genotypic variation, molecular phenotype variation, and phenotypic variation for a particular complex trait^46–50^. Notwithstanding recent methodological concerns^51^, the TWAS approach can provide meaningful insights into the molecular basis of quantitative trait loci and an integrated knowledgebase of tissue-specific human TWAS associations, the TWAS Atlas, has recently been developed^52^. TWAS approaches have also been leveraged to identify genes with expression patterns that modulate phenotypic variability for economically important traits in cattle^53,54^. Various TWAS methods have been developed to study the effects of proximal genetic variants (*cis*-eQTLs) on transcriptional regulation^46,47,55,56^ that do not consider distal/interchromosomal regulatory polymorphisms (*trans*-eQTLs), which are a major component of the omnigenic model of complex trait inheritance^57^. To address this, the Multi-Omic Strategies for TWAS (MOSTWAS) suite of tools has been developed, which extend traditional TWAS approaches to include *trans*-acting variants around regulatory biomarkers (e.g., transcription factor and microRNA genes) to increase the power to detect significant gene-trait associations^58^.

It has previously been reported that the peripheral blood (PB) immune responses reflect those at the site of infection for bTB disease^59^. In this regard, our group and others have detected and characterised PB transcriptional biosignatures of *M. bovis* infection and bTB^60–70^. However, functional integration of PB transcriptomes, host genomic variation, and GWAS data sets for bTB susceptibility has not been performed previously. Additionally, to-date there have been no published studies that use the TWAS approach to understand the regulatory genome in the context of the host response to mycobacterial infections that cause TB in mammals. Therefore, using PB RNA-seq data from *M. bovis*-infected and control non-infected cattle, and imputed genome-wide SNP data, we combine an eQTL analysis with multiple bTB GWAS data sets^33^ and conduct a summary TWAS incorporating *trans*-acting genomic variants^58^, which identifies important new genes underpinning the mammalian host response to mycobacterial infections that cause TB.

## Results

### Animal disease phenotyping

**Fig. 1** provides an overview of the experimental workflow and computational pipeline used for this study. A large panel of bTB reactor (bTB+; *n* = 60) and control (bTB−; *n* = 63) cattle were recruited that had a positive (reactor) and negative reaction, respectively, to the single intradermal comparative tuberculin test (SICTT). All animals were male, and the mean age of the animals was 21.9 ± 8.3 months. **Supplementary Table 1** provides detailed information about these animals, including the last four digits of the ear tag ID, date of sampling, and breed ancestry based on comprehensive pedigree information.

**Fig. 1:**
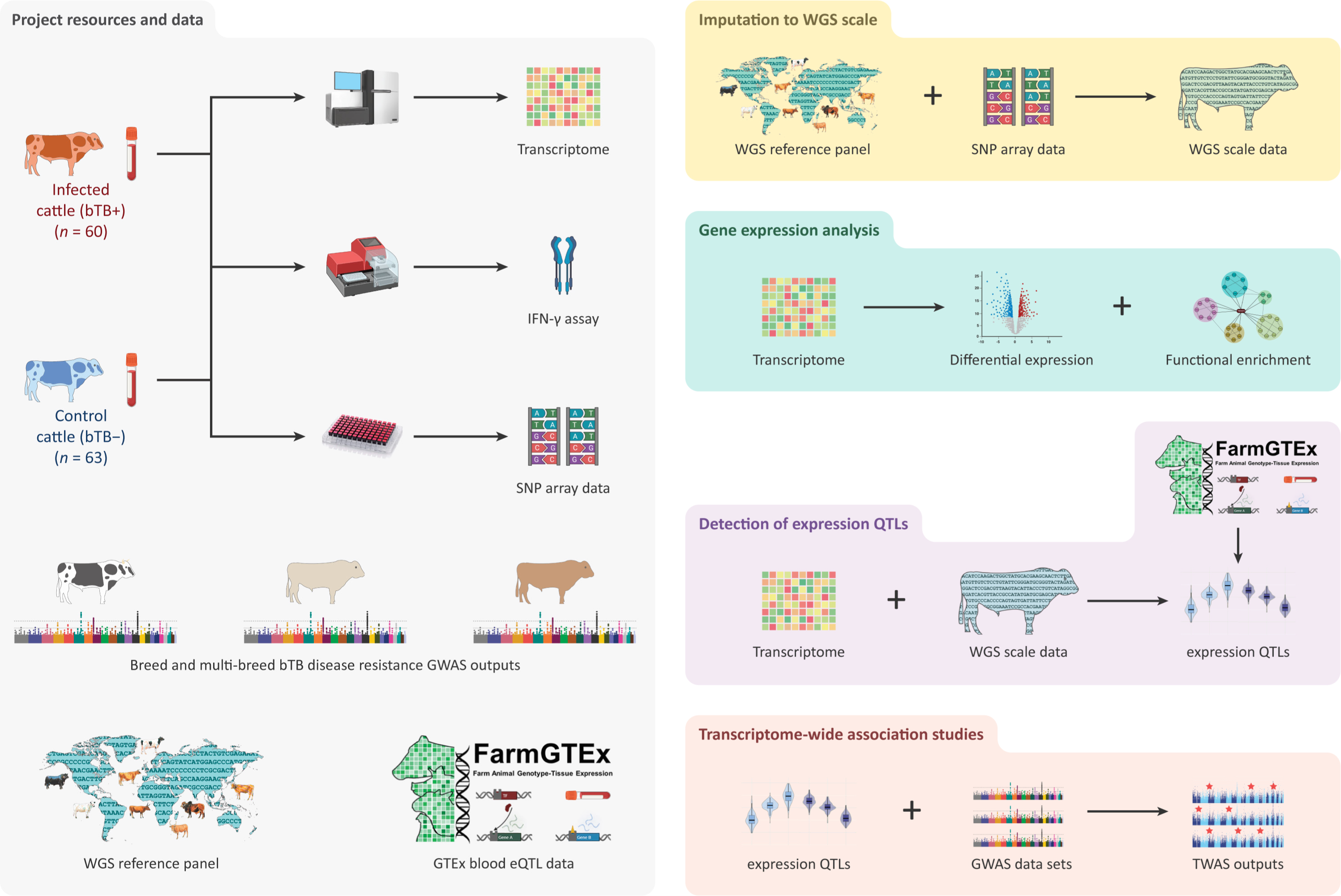
Experimental and computational workflow. Data resources for the project included; 1) newly generated high-resolution SNP-array data, peripheral blood RNA-seq data and interferon gamma (IFN-γ) release assay (IGRA) measurements from a reference panel of *n* = 60 bovine tuberculosis (bTB) reactor (bTB+) and *n =* 63 control (bTB−) cattle; 2) single and multi-breed GWAS summary statistics for bTB susceptibility from Ring *et al*., (2019)^33^; 3) whole genome sequence (WGS) data from Dutta *et al*., (2020)^72^ and; 4) whole blood eQTL summary statistics from the Cattle GTEx consortium^53^. For the reference panel, SNP array genotype data was remapped to the ARS-UCD1.2 bovine genome build and imputed using the WGS cohort as a reference panel. RNA-seq data was aligned to ARS-UCD1.2 with the resulting count matrices normalised using various methodologies (See Methods) for inclusion in the differential expression, functional enrichment, and expression quantitative trait loci (eQTL) analyses. The normalised expression matrix was integrated with the imputed SNP-array data for the eQTL analysis. To assess the replication of eQTLs, we leveraged whole blood eQTL summary statistics from the Cattle GTEx consortium^53^ and separately performed various permutation tests on identified *trans*-eQTLs. Finally, the GWAS summary statistics were remapped to ARS-UCD1.2 before being integrated with the reference panel eQTL results to conduct three single- and one multi-breed transcriptome wide association study (TWAS) for bTB susceptibility using the MOSTWAS software^58^ (some figure components created with a Biorender.com license).

For the purposes of this study, and as a confirmatory test, the interferon gamma (IFN-γ) diagnostic assay was used to evaluate *M. bovis* infection status in all 123 recruited animals. The criterion for IFN-γ test positivity was a test result difference greater than 80 ELISA units for the purified protein derivative (PPD)-bovine (PPDb) IFN-γ value minus the PPD-avian (PPDa) IFN-γ value (ΔPPD)^71^. The mean ΔPPD (± SE) for the bTB+ animal group was 1170.35 ± 84.48 compared to −360.46 ± 55.17 for the bTB− group and this group difference was highly significant (two tailed Wilcoxon rank-sum test; *P* < 3.258 × 10^−21^) (**Supplementary Fig. 1, Supplementary Table 2**). One designated bTB− control animal produced a positive result for the IFN-γ test (C050, ΔPPD = 263.1) and two designated bTB+ animals elicited a negative result (T007, ΔPPD = 36.0; T062, ΔPPD = −52.3). These results yielded test sensitivity and specificity rates of 96.67% and 98.41%, respectively, which is in line with IFN-γ test performance under Irish conditions^71^. These animals were still designated as bTB− and bTB+, respectively, and included in subsequent analyses.

### RNA-seq mapping statistics and genome-wide SNP imputation

Peripheral blood RNA sequencing yielded a mean of 35,129,315 ± 3,430,729 reads per individual sample library (*n* = 123 libraries and ± standard deviation). Reads were aligned to the ARS-UCD1.2 *B. taurus* genome build with a mean of 33,352,903 ± 3,206,593 (95.06% ± 0.76%) reads mapping uniquely, 779,866 ± 110,837 (2.22% ± 0.17%) mapping to multiple loci, 14,168 ± 2,639 (0.04 ± 0.008%) mapping to an excessive number of loci, 97,358 ± 274,880 (2.74 ± 0.68%) that were too short, and 15,020 ± 2,867 (0.04% ± 0.008%) that could not be assigned to any genomic locus. The mean mapped length was 297.8 ± 0.3 bp (**Supplementary Table 3**). None of the libraries exhibited an abnormal distribution of gene counts (**Supplementary Fig. 2**).

A total of 591,947 array-genotyped SNPs were available for analysis. To determine if any animal samples were inadvertently duplicated, we first LD-pruned the array genotype data following filtering of variants which were rare (MAF < 0.1) and which deviated from HWE (*P* < 1 × 10^-6^) to yield 34,272 SNPs. We then calculated the identity by state (IBS) among all animals using PLINK. We set a cut-off of 0.85 for deeming two samples as duplicates. All pairs of animals returned an IBS distance value < 0.8 (**Supplementary Fig. 3a**). Following this, we remapped the raw SNP array-data from the UMD 3.1 genome build to the ARS-UCD 1.2 reference genome and imputed the remapped variants up to WGS scale using a Global Reference Panel as a reference^72^ (**Supplementary Note 4**). Imputation performance increased as MAF increased with poor performance observed at variants with a MAF ≤ 1% (**Supplementary Fig. 5)**. Following the removal of imputed variants that displayed poor imputation performance (R^2^ < 0.6), possessed a low MAF (< 0.05), and that deviated significantly from HWE (*P <* 1 × 10^-6^), a total of 3,866,506 imputed autosomal SNPs were retained for the eQTL analysis. Lastly, comparison of the imputed SNP profiles with RNA-seq reads using QTLtools showed that there were no sample mismatches and that the imputed WGS data correctly matched the transcriptomics data for all animals (**Supplementary Fig. 3b**).

### Population genomics, differential gene expression, and functional enrichment analyses

The results of the genetic structure analysis using the ADMIXTURE program with 34,272 pruned genome-wide array SNPs and an inferred number of ancestral populations *K* = 2 are shown in **Fig. 2a**. A principal component analysis (PCA) plot of principal components (PC) 1 and 2 generated from the same set of pruned SNPs is shown in **Fig. 2b** with percentage Holstein ancestry and component 1 from the ADMIXTURE structure analysis also shown for each animal sample (see also **Supplementary Table 4**). The results of these two analyses were mutually compatible; component 1 from the ADMIXTURE structure plot was in concordance with PC1 (10.8% of the variance derived from the top 20 PCs from the PCA and likely corresponded, at least in part, to Holstein ancestry for the animals that had pedigree-derived breed composition data (113 out of 123 animals) (**Fig. 2b**, **Supplementary Table 1**). There was also a highly significant positive correlation (Spearman correlation (*ρ*) = 0.829, *P* < 2.2 × 10^-16^) between the pedigree-derived percentage Holstein ancestry values and component 1 from the ADMIXTURE structure plot (**Supplementary Fig. 6a**). PC2 (8.0% of the total variance of the top 20 PCs) likely accounts for population structure within the Holstein-Friesian populations, which has been documented previously in an independent cohort^27^. We observed that the genetic structure of the study population (bTB− and bTB+) was a confounder in the transcriptomics data set because there was sample clustering caused by breed ancestry observed in the PCA of the top 1,500 most variable genes determined from the variance stabilised transformed (VST) count matrix in DESeq2 (**Supplementary Fig. 6b**).

**Fig. 2:**
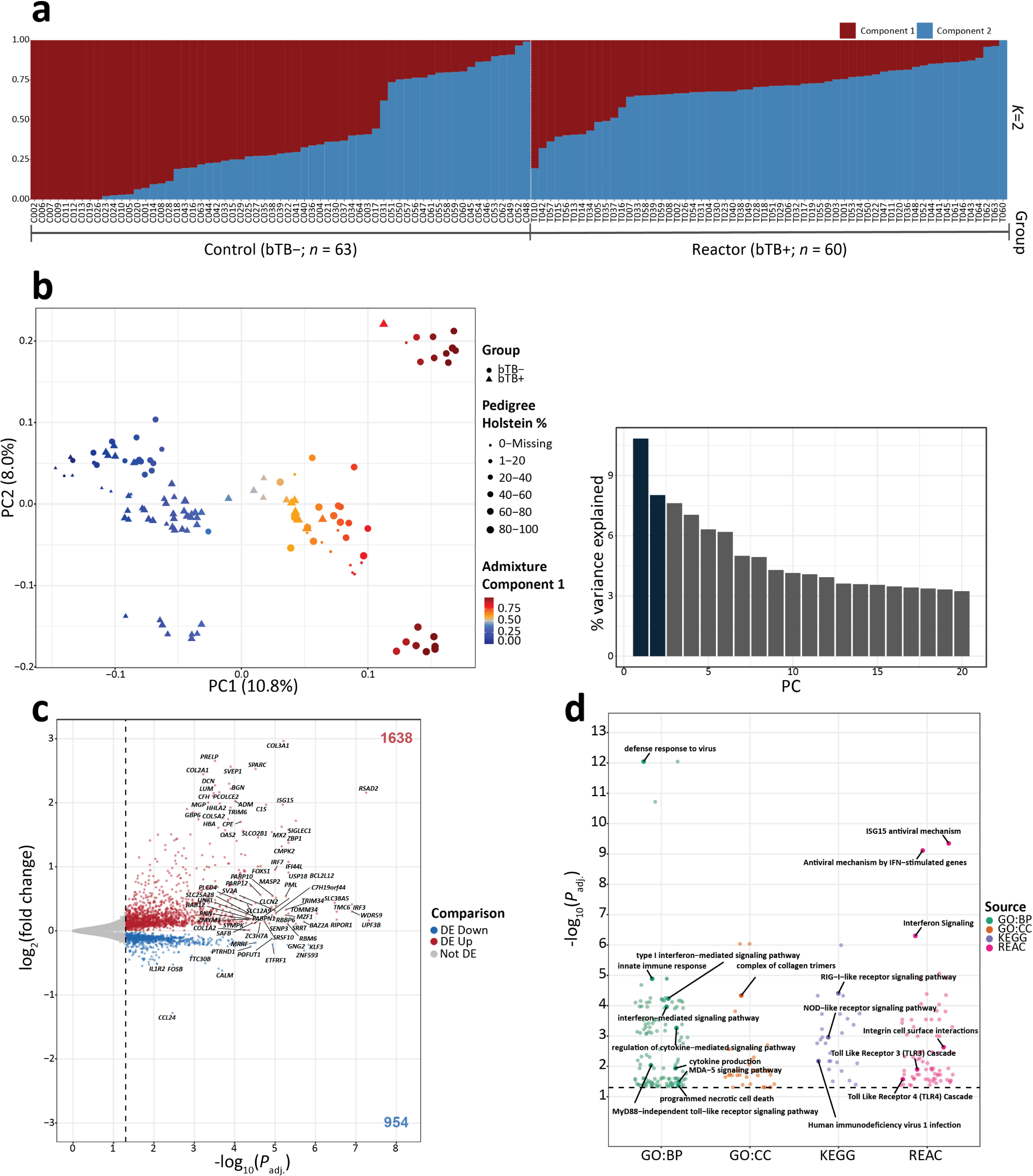
Population genomics, differential expression, and functional enrichment analyses. **a** Structure plot showing the proportion of ancestry components 1 and 2 from the ADMIXTURE analysis for 123 animals (reactor bTB+ and control bTB−). **b** Principal component analysis (PCA) plot of PC1 and PC2 derived from 34,272 pruned SNPs genotyped in 123 animals. The data points are shaped based on their experimental designation, coloured based on the inferred ancestry component 1 from the ADMIXTURE analysis, and sized based on their reported pedigree Holstein percentage. A histogram plot of the relative variance contributions for the first 20 PCs is also shown with PC1 and PC2 accounting for 10.8% and 8.0% of the variation in the top 20 PCs, respectively. **c** Horizontal volcano plot of differentially expressed genes (DEGs) for the bTB+ (*n* = 60) versus bTB− (*n* = 63) contrast with thresholds determined by FDR *P*_adj._ < 0.05 and an absolute log_2_ fold-change (LFC) > 0. The *x*-axis represents the -log_10_ *P*_adj._ and the *y*-axis represents the log_2_ fold change. **d** Jitter plot of significantly impacted pathways/GO terms identified across the Gene Ontology (GO) Biological Processes (GO:BP), Cellular Compartment (GO:CC), Reactome (REAC) and Kyoto Encyclopaedia of Genes and Genomes (KEGG) databases using g:Profiler. The data points are coloured according to the corresponding database.

We performed a differential expression analysis (DEA) to identify differentially expressed genes (DEGs) between the reactor (bTB+) and control (bTB−) animal groups, which incorporated PC1 and PC2 (**Fig. 2b**), age in months, and sequencing batch (1 or 2) as covariates in the generalised linear model. With this approach, we identified 2,592 DEGs (FDR *P*_adj._ < 0.05) for the bTB+ versus bTB− contrast (**Fig. 2c**, **Supplementary Table 5**). Within the bTB+ group, increased expression was observed for 1,638 DEGs and 954 DEGs exhibited decreased expression. We then selected a subset of 1,091 highly significant DEGs (FDR *P*_adj._ < 0.01) for gene set overrepresentation and functional enrichment analyses using g:Profiler and IPA^®^. In this subset of DEGs, 602 and 489 genes exhibited increased and decreased expression, respectively in the bTB+ cohort.

Using the g:Profiler tool (FDR *P*_adj._ < 0.05) we observed a clear enrichment for innate immune response, pathogen internalisation, and host-pathogen interaction GO terms and biological pathways (**Fig. 2d**). The top significantly enriched functional entity was the *Defense response to virus* (FDR *P*_adj._ = 9.02 × 10^-13^) GO:BP term. Other significantly enriched functional entities included: *Cytosolic pattern recognition receptor signalling pathway* (FDR *P*_adj._ = 1.24 × 10^-4^) GO:BP term; *RIG-I-like receptor signalling pathway* (FDR *P*_adj._ = 3.88 × 10^-5^) from KEGG; and *Antiviral mechanism by IFN-stimulated genes* (FDR *P*_adj._ = 7.67 × 10^-10^) from the Reactome database. All significant results obtained from g:Profiler, in addition to the intersection of DE genes with the respective functional entities, are provided in **Supplementary Table 6**. For IPA^®^, a total of 996 DE genes and 14,228 background genes were successfully mapped. The significantly enriched (FDR *P*_adj._ < 0.05) pathways identified from IPA^®^ included *Interferon alpha/beta signalling* (FDR *P*_adj._ = 4.09 × 10^-8^), *Oxidative phosphorylation* (FDR *P*_adj._ = 2.91 × 10^-6^) and *Activation of IRF by cytosolic pattern recognition receptors* (FDR *P*_adj._ = 9.12 × 10^-3^) (**Supplementary Fig. 7a**, **Supplementary Table 7**). Upstream transcriptional regulator analysis using IPA^®^ revealed that the transcriptional regulator, ETV3 was the most significant upstream biological regulator of the inputted DE genes (FDR *P*_adj._ = 4.89 × 10^-19^) (**Supplementary Fig. 7b**) Other important statistically significant (FDR *P*_adj._ < 0.05) upstream regulators include TLR3, STING1, IRF5, and STAT1 (for complete results see **Supplementary Table 8**).

### Identification of *cis*-expression quantitative trait loci

We used a linear regression model in TensorQTL to test associations between expressed genes and SNPs that passed filtering thresholds to identify local (± 1 Mb) *cis*-eQTLs in the reactor (bTB+) group (*n* = 60), the control (bTB−) group (*n =* 63), and a combined all animals group (AAG, *n* = 123). As covariates, we also included 1) the top five SNP genetic variation PCs (PC1-5) inferred for each group separately to account for interbreed differences between the animals; 2) age in months; 3) sequencing batch; 4) disease status (where applicable); and 5) transcriptomic PCs with PC1-8, PC1-9, and PC1-14 for the bTB+, bTB−, and AAG cohorts, respectively. The number of transcriptomic PCs to use was determined using the elbow method (**Supplementary Fig. 8**). We also removed known covariates (genotype PCs, age, batch, disease status) that were well captured by the inferred covariates (unadjusted *R*^2^ ≥ 0.9) and the final set of covariates for each cohort are detailed in **Supplementary Tables 9-11**. In total, we tested 14,701, 14,598, and 14,612 genes in the bTB+, bTB−, and AAG cohorts, respectively for *cis* SNP variants associated with their expression levels (**Supplementary Fig. 9a**).

**Table 1** summarises the number of significant (FDR *P*_adj._ < 0.05) *cis*-eQTLs, *cis*-eVariants, and *cis-*eGenes identified in all three groups. We identified 2,235, 3,419, and 6,676 *cis*-eGenes in the bTB+, bTB−, and AAG cohorts, respectively, with the largest proportion captured by the AAG group (**Fig. 3a, Supplementary Tables 12-14**). For each *cis*-eGene in each group, variants with a nominal *P*-value below the gene-level threshold (**Supplementary Fig. 9b**) were considered significant *cis*-eVariants. Overall, we identified 168,251, 415,861 and 1,103,004 significant *cis*-eVariant:gene associations in the bTB+, bTB−, and AAG cohorts. Of these *cis*-eVariants, 21.0%, 23.1% and 35.7% were associated with >1 *cis*-eGene. For all three groups, we identified hundreds to thousands of *cis*-eGenes with multiple independent acting *cis*-eQTLs (**Fig. 3b**). The conditional analysis detected 13.2%, 27.1% and 80.1% additional independent *cis*-eQTLs in the bTB+, bTB−, and AAG cohorts, respectively. We observed that the top *cis*-eQTL identified by the permutation analysis tended to cluster close to the transcriptional start site (TSS) of the associated gene, whereas conditionally independent *cis-*eQTLs were located at variable distances to the TSS (Wilcoxon rank-sum test; *P* < 2.2 × 10^-16^) (**Fig. 3c**). The permuted and conditional *cis*-eQTL associations were symmetrical around the TSS with no enrichment in the 5’ or 3’ directions (**Supplementary Fig. 10**). We noted a moderately negative but highly significant Spearman correlation in the effect size estimates of *cis*-eQTLs and their respective distances to the TSS of the associated gene in all three groups (**Fig. 3d**).

**Table 1:** Total number of and unique number of significant (FDR *P*_adj._ < 0.05) *cis*-eQTLs, *cis-*eVariants and their corresponding *cis*-eGenes identified across the reactor (bTB+), control (bTB−) and combined all animals (AAG) cohorts, respectively.

| Class of eQTL/eGene | Distance between associated pair | bTB+ (n = 60) | bTB– (n = 63) | AAG (n = 123) |
| --- | --- | --- | --- | --- |
| *cis*-eQTLs (permutation) | ± 1 Mb | 2,235 | 3,419 | 6,676 |
| Conditionally independent *cis*-eQTLs | ± 1 Mb | 295 | 925 | 5,385 |
| Total number of independent *cis*-eQTLs | ± 1 Mb | 2,530 | 4,344 | 12,061 |
| *cis*-eGenes | ± 1 Mb | 2,235 | 3,419 | 6,676 |
| *cis*-eVariant associations | ± 1 Mb | 168,251 | 415,861 | 1,103,004 |
| Unique *cis*-eVariants | ± 1 Mb | 139,616 | 319,734 | 709,337 |

**Fig. 3:**
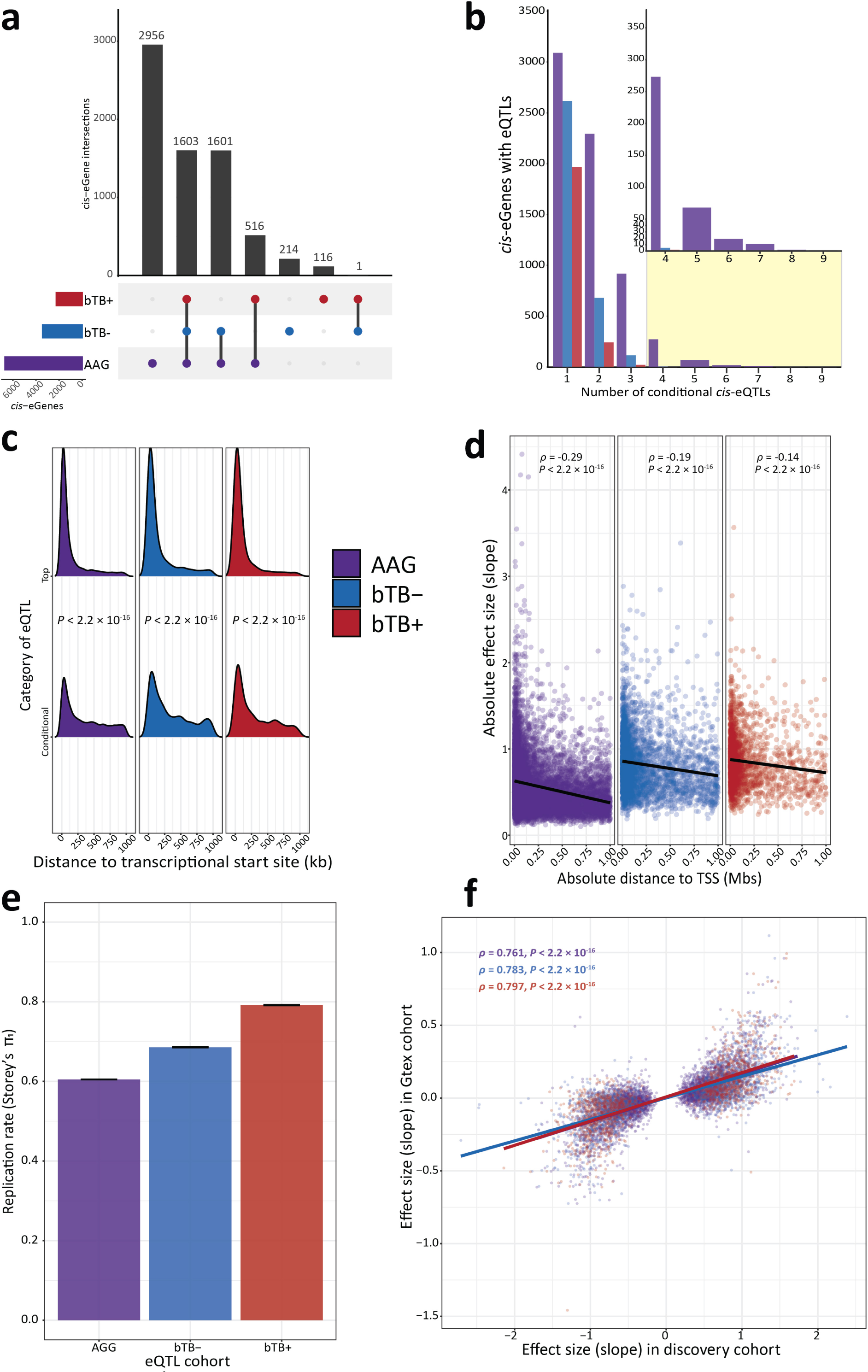
*Cis*-expression quantitative trait loci (eQTL) mapping and external replication results. **a** Upset plot showing the intersection of shared *cis*-eGenes identified in the reactor (bTB+), control (bTB−), and combined all animal (AAG) cohorts, respectively. **b** Barplot illustrating the number of genes with a significant primary or conditional *cis*-eQTL for degrees 1–9 across all three groups. Inset shows the number of genes for *cis*-eQTL degrees 4–9. **c** Ridgeline plot showing the distribution of the absolute distance from the transcriptional start site (TSS) of top and conditional *cis*-eQTLs identified in all three groups. *P*-values are inferred from the Wilcoxon rank-sum test between top and conditional *cis*-eQTLs within each group. **d** Scatter plot illustrating the relationship between absolute *cis*-eQTL effect size and distance to the TSS for all significant *cis*-eQTLs (top and conditional) identified in each group separately. Spearman correlation values are also reported in addition to the corresponding *P*-value representing the significance level of each respective correlation. The black line indicates line of best fit. **e** Replication rate as measured by Storey’s π_1_ in the current study and whole blood *cis*-eQTLs identified in the Cattle GTEx^53^. The error bars indicate the standard error from 100 bootstrap samplings. **f** Scatterplot illustrating the effect sizes of significant *cis*-eQTLs identified in this study and matched variant-gene pairs identified in the Cattle GTEx. Spearman correlation values are also reported in addition to the corresponding *P*-value representing the significance level of each respective correlation. The coloured lines indicate lines of best fit within each group, respectively. The colour scheme for each group is consistent throughout the figure.

To assess replication of the *cis*-eQTLs identified in this study, we used three metrics: allelic concordance (AC), the π_1_ statistic to measure the proportion of true positive associations, and the Spearman correlation coefficient of effect size estimates in an external set of whole-blood *cis*-eQTL summary statistics obtained from the Cattle GTEx Consortium^53^. We observed high AC between top and significant *cis*-eQTLs identified in this study and significant *cis*-eQTLs identified in the Cattle GTEx (AC_bTB+_ = 99.27%, AC_bTB−_ = 99.10%, and AC_AAG_ = 98.87%). We observed moderate to high π_1_ statistics across all groups indicating good replication (π_1bTB+_ = 0.791 ± 0.0009, π_1bTB−_ = 0.685 ± 0.0007, and π_1AAG_ = 0.605 ± 0.0004) (**Fig. 3e**). We also noted a positive and significant Spearman correlation in effect size estimates for the top significant eQTLs identified in our study and the matched variants from the Cattle GTEx across all three groups (*ρ*_bTB+_ = 0.797, *ρ*_bTB−_ = 0.783, and *ρ*_AAG_ = 0.761) (**Fig. 3f**).

### *Cis*-eQTLs regulate the peripheral blood transcriptional response to *M. bovis* infection

We compared the *cis*-eGenes identified in the bTB+ and the bTB− groups and that were also replicated in the AAG group (**Fig. 3a**) to assess if there were genomic variants influencing the PB transcriptomes for each of these biological states. This approach facilitated identification of; 509 bTB+ only (bTB+) *cis*-eGenes, identified in the bTB+ and AAG cohorts but not the bTB− group; 1603 *cis*-eGenes that were identified across all three groups; and 1593 bTB− only (bTB−) *cis*-eGenes identified in the bTB− and AAG cohorts but not the bTB+ group (**Fig. 4a**). We then performed a gene set overrepresentation analysis of *cis*-eGenes for four groups (bTB−, bTB− and AAG, bTB+, bTB+ and AAG) using g:Profiler (**Fig. 4b**). Significantly overrepresented (FDR *P*_adj._ < 0.05) functional entities identified using *cis*-eGenes identified in the bTB− only and the AAG cohorts included the *MHC class II protein complex* GO:CC term; The *ER-Phagosome pathway* Reactome term and the *Leishmaniasis* KEGG term. Considering *cis*-eGenes identified only in the bTB− group, significantly impacted pathways included the *Succinyl-CoA metabolic process* and the *antigen processing and presentation of peptide antigen* GO:BP terms*. Cis*-eGenes identified in the bTB+ and AAG group were significantly overrepresented in pathways that included the *Th1 and Th2* and *Th17 cell differentiation* KEGG terms and *Phsophorylation of CD3 and TCR Zeta chains* Reactome term. In the bTB+ group, we observed a number of GO:BP terms significantly overrepresented by *cis*-eGenes including; *Negative regulation of chemokine (C-C motif) ligand 4* and *5 production* and *Negative regulation of macrophage inflammatory protein 1 alpha production*. All overrepresentation results obtained using g:Profiler for the analysis of these four gene sets are detailed in **Supplementary Table 15-18**.

**Fig. 4:**
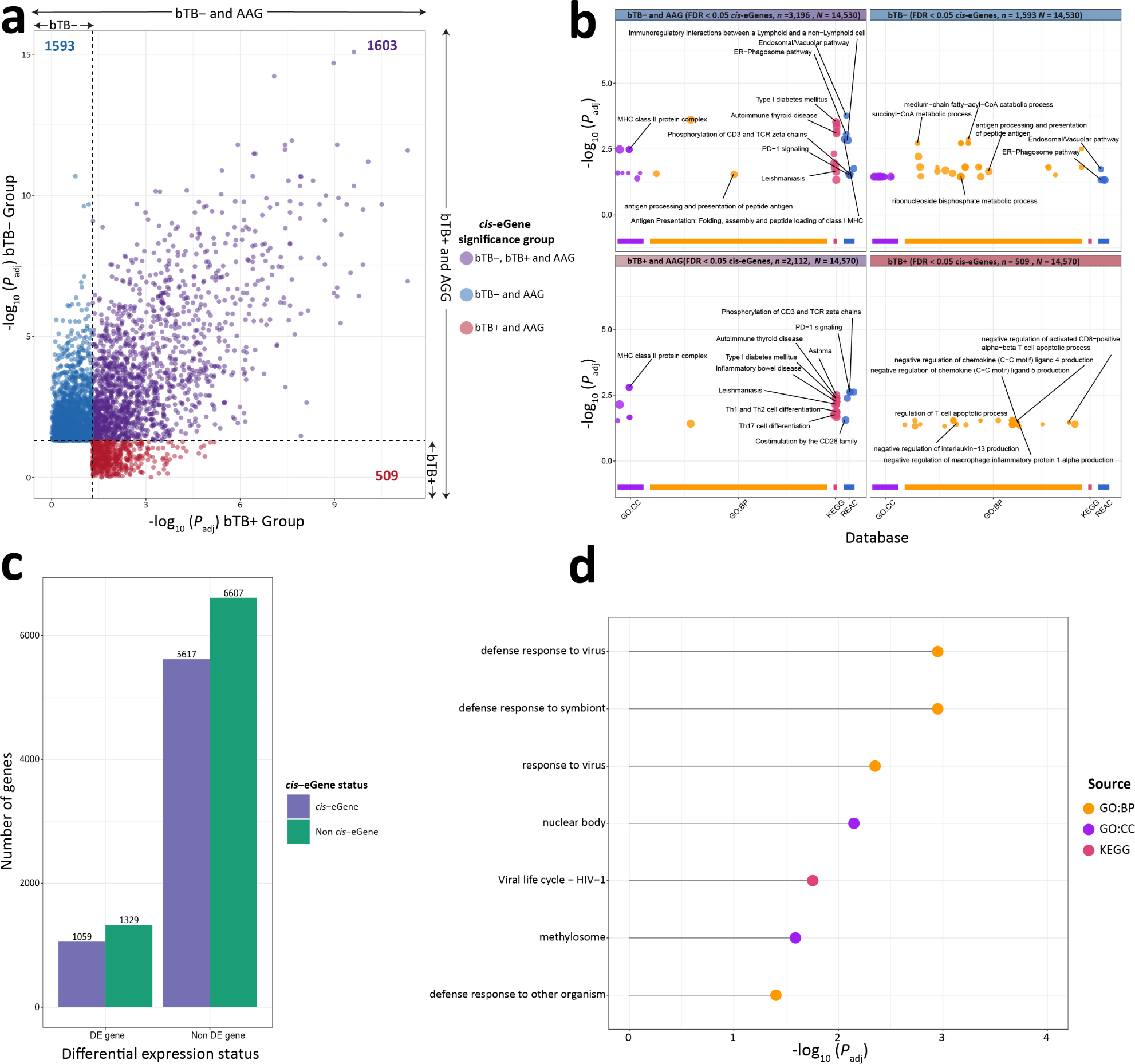
Integrative *cis*-eGene analysis. **a** Scatter plot of significant *cis*-eGenes identified in the control (bTB−) and combined all animals (AAG) cohorts but not the reactor (bTB+); those identified across all three groups (bTB−, bTB+, and AAG); and those identified in the bTB+ and AAG cohorts but not the bTB− group (bTB+). The *y*-axis corresponds to the most significant FDR *P*_adj._ variant-gene pair identified in the bTB− group and the *x*-axis corresponds to the most significant *P*_adj._ variant-gene pair identified in the bTB+ group. Dashed lines indicate and FDR *P*_adj._ < 0.05. **b** Significantly impacted pathways/GO terms by *cis*-eGenes from the bTB−, bTB−/AAG, bTB+, and bTB+/AAG cohorts across Gene Ontology (GO) Biological Processes (GO:BP), Cellular Compartment (GO:CC), Reactome (REAC), and the Kyoto Encyclopaedia of Genes and Genomes (KEGG) databases using g:Profiler. The number of input genes for each set (*n*) and number of background genes (*N*) for each set is also detailed. Data points are coloured based on their corresponding database **c** Barplot showing the classification of genes tested in both the *cis*-eQTL and differential expression analysis that were classified as differentially expressed (DE) or not DE *cis*-eGenes. **d** Lollipop chart showing significantly impacted pathways and GO terms (FDR *P*_adj._ *<* 0.05) for the 1,059 DE-*cis*-eGenes across the GO:BP, GO:CC, and KEGG databases. The pathways are ordered based on adjusted *P*-value and are coloured based on their corresponding database.

To identify DE *cis*-eGenes, we focused on the *cis*-Genes identified in the AAG group that overlapped with the DEG results (**Fig. 2c**). Of the 2,592 DE genes, 2,388 (92.12%) were tested in the *cis-*eQTL analysis. A total of 1,059 DEGs were characterised as *cis*-eGenes and 1,329 DEGs were not (**Fig. 4c**). We did not identify a significant association between DEGs and genes characterised as being *cis*-eGenes (chi-square test; χ^2^ = 2.0068, *P* = 0.1566). For the 1,059 DE *cis-*eGenes, we conducted a g:Profiler overrepresentation analysis using the set of genes that overlapped between the DEG and the AAG *cis*-eQTL analyses as the background set. Significantly impacted pathways and GO terms perturbed by these DE *cis-*eGenes included the *Defense response to virus* GO:BP term (FDR *P*_adj._ = 1.12 × 10^-3^), the *Viral life cycle – HIV-1* (FDR *P*_adj._ = 1.76 × 10^-2^) KEGG pathway and the *Methylosome* (FDR *P*_adj._ = 2.57 × 10^-2^) GO:CC term (**Fig. 4d**, **Supplementary Table 19**).

### Mapping of *trans*-expression quantitative trait loci is confounded by bovine population genetic structure

We employed a linear regression model in QTLtools that included the same inputs as the *cis*-eQTL mapping procedure to characterise distal intrachromosomal (> 5 Mb) and interchromosomal *trans*-eVariants. **Table 2** summarizes the numbers of intra- and interchromosomal *trans*-eVariants and *trans*-eGenes detected in all three groups (bTB+, bTB−, and AAG) and **Fig. 5a** shows the overlap of *trans*-eGenes across these groups. In total, we identified 497, 916, and 5,314 *trans*-eVariants (FDR *P*_adj._ < 0.05) in the bTB+, bTB−, and AAG cohorts, which were associated with 13, 17 and 107 *trans*-eGenes, respectively (**Fig. 5a, Supplementary Table 20-22**). Because of the relatively small numbers of *trans*-eGenes identified in the bTB+ and bTB− groups, we focused on the AAG set of *trans*-eGenes for a more detailed analysis.

**Table 2:** Total number of and unique number of significant (FDR *P*_adj._ < 0.05) intrachromosomal and interchromosomal *trans-*eVariants and *trans*-eGenes identified across the reactor (bTB+), control (bTB−) and combined all animals (AAG) cohorts, respectively.

| Class of eQTL/eGene | Distance between associated pair | bTB+ (n = 60) | bTB- (n = 63) | AAG (n = 123) |
| --- | --- | --- | --- | --- |
| *Trans*-eVariant associations | > 5 Mb | 497 | 916 | 5,314 |
| *Trans*-eGenes | > 5 Mb | 13 | 17 | 107 |
| Unique *trans*-eVariants | > 5 Mb | 497 | 704 | 4,976 |
| Intrachromosomal *trans*-eVariants | > 5 Mb and on same chromosome | 0 | 215 | 588 |
| Interchromosomal *trans*-eVariants | Different chromosome | 497 | 701 | 4,726 |
| Intrachromosomal *trans*-eGenes | > 5Mb and on same chromosome | 0 | 2 | 26 |
| Interchromosomal *trans*-eGenes | Different chromosome | 13 | 15 | 81 |

**Fig. 5:**
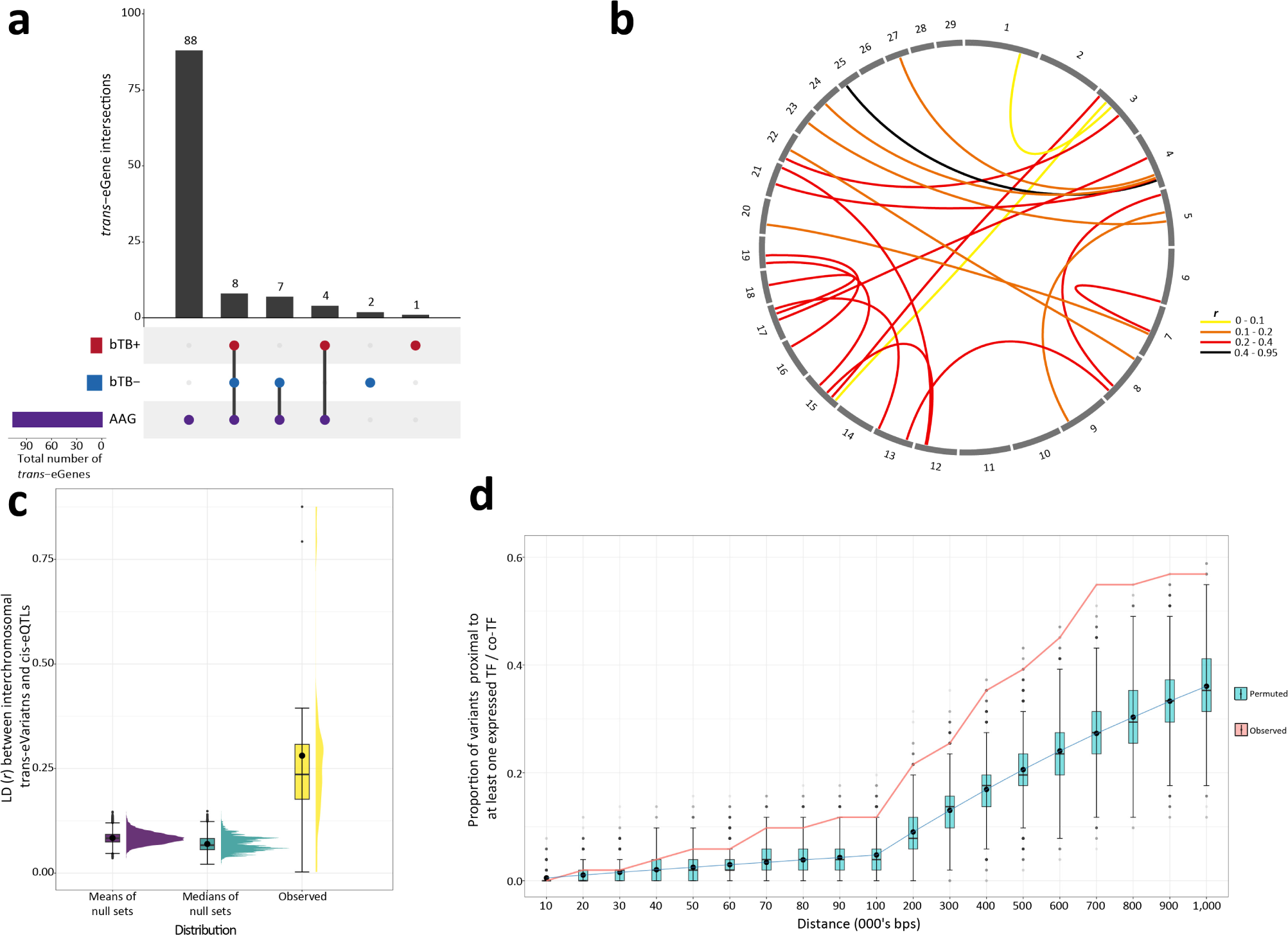
*Trans*-expression quantitative loci mapping results and downstream analysis of inter-chromosomal *trans*-eVariants. **a** Upset plot showing the intersection of shared *trans*-eGenes identified in the reactor (bTB+), the control (bTB−), and the combined all animal group (AAG) cohorts, respectively. **b** Circos plot showing the linkage disequilibrium (LD) (*r*) relationship between the top interchromosomal *trans*-eVariants and top *cis*-eQTLs for the same gene. **c** Comparison of observed interchromosomal LD relationship (*r*) between 23 top interchromosomal *trans*-eVariants and *cis*-eQTLs of the same gene versus the mean and median distributions of 10,000 sets of 23 interchromosomal null *trans*-eVariant and top *cis*-eQTL pairs. **d** The proportion of observed (blue) highly significant top *trans*-eVariants (FDR *P*_adj._ < 0.01) not in LD with top *cis*-eQTLs of the same gene (*r*^2^ < 0.01) residing close to at least one expressed transcription factor (TF) or TF-cofactor at various (±) intervals versus 10,000 null sets of 51 SNPs (red). Horizontal lines inside the boxplots show the medians, solid circles indicate the means. Box bounds show the lower quartile (Q1, the 25^th^ percentile) and the upper quartile (Q3, the 75^th^ percentile). Whiskers are minima (Q1 – 1.5 × IQR) and maxima (Q3 + 1.5 × IQR) where IQR is the interquartile range (Q3-Q1).

Identification, biological interpretation, and replication of peripheral blood *trans*-eQTLs is challenging owing to the heterogenous nature of the tissue and the small effect sizes associated with putative distal variants^73^; however, notwithstanding these limitations, we observed an inflated number of *trans*-eQTLs in this study compared to previous reports in humans^74^. We first focused on the 588 intrachromosomal *trans*-eVariants associated with 26 *trans*-eGenes (**Supplementary Fig. 11a**). Among the intrachromosomal *trans*-eVariants associated with the same intrachromosomal *trans*-eGene, we observed a high LD genotype correlation between these variants (*r* = 0.889 ± 0.243SD) (**Supplementary Fig. 11b**). We therefore selected the most significant intrachromosomal *trans*-eVariant for each gene and computed the LD between this variant and the top *cis*-eQTL of the same gene. In total, 24 genes had a significant *cis*-eQTL and intrachromosomal *trans*-eVariant associated with its expression levels. We observed high LD amongst top intrachromosomal *trans*-eVariants and top *cis*-eQTLs of the same gene (*r* = 0.562 ± 0.203SD). To determine whether our observed LD pattern was significantly greater than what would be expected by chance, we randomly sampled 10,000 sets of 24 variant pairs which were no less than 5 Mb and no greater than 14,065,301 bp apart (the latter cutoff was two standard deviations of the distribution of distances between top *trans*-eVariants and top *cis*-eQTLs for the same gene). We calculated the medians and means of these 10,000 sets to generate two null distributions. We then calculated a permuted *P*-value (*P*_perm._) defined as the proportion of permutations with a median and mean intrachromosomal LD relationship at least as large or greater than the observed set. After this procedure, we obtained a permuted *P*-value of < 0.0001 indicating that our observed set of intrachromosomal *trans*-eVariants was significantly inflated by LD (**Supplementary** Fig. 11c**, Supplementary Table 23**).

We next focused on the 4,726 interchromosomal *trans*-eVariants associated with 81 *trans*-eGenes. We again selected the most significant SNP associated with each *trans*-eGene and calculated the LD between these interchromosomal *trans*-eVariants and the top *cis*-eQTL of the same gene. In total, 23 genes had a significant interchromosomal *trans*-eVariant and *cis*-eQTL associated with its expression levels. We observed a complex interchromosomal LD pattern between *cis*-eVariants and *trans*-eVariants of the same gene (*r* = 0.280 ± 0.199SD) (**Fig. 5b**). To assess whether our observed LD pattern was significantly greater than what would be expected by chance, we first sampled for each *trans*-eVariant with replacement, 1000 null *trans*-eVariants from the same chromosome and same allele frequency as putative *trans*-eVariants. We then computed the LD relationship between these null *trans*-eVariants and the *cis*-eQTLs of interest and then randomly generated 10,000 sets of 23 null interchromosomal *trans*-eVariants and the corresponding top *cis*-eQTL pairs. We performed the same procedure used for the intrachromosomal analysis to generate two null distributions with two *P*_perm._ values < 0.0001, which indicated that our top interchomosomal *trans*-eVariants were in high LD with top *cis*-eQTLs of the same gene (**Fig. 5c**, **Supplementary Table 24**).

### *Trans*-eVariants cluster close to expressed transcription factors and co-transcription factors

We next filtered putative *trans*-eVariants to retain variants with a highly significant (FDR *P*_adj._

< 0.01) *trans* association and that were not in LD (genotype squared correlation (*r*^2^) > 0.01) with a *cis*-eQTL of the same gene. This reduced the number of *trans*-eVariants and *trans*-eGenes available for analysis to 3,934 and 51, respectively. We hypothesised that these *trans*-acting variants resided close to expressed transcription factors (e-TFs) or transcription factor co-factors (e-coTFs) and would regulate *trans*-eGenes through influencing expression of the e-TFs/e-coTFs in *cis*. To investigate this, we first selected the top *trans-*eVariant associated with each *trans*-eGene and downloaded the genomic locations of 2,384 annotated TFs/coTFs from the AnimalTFDB v.4.0 database^75^. Of these, 973 (40.81%) passed expression filtering thresholds for inclusion in the AAG eQTL analysis. We next calculated the proportion of the 51 most significant *trans*-eVariants that resided close to at least one of the 973 TFs/co-TFs at various distances ranging from ±10 kb to ±1 Mb versus a random set of 51 SNPs computed 10,000 times to generate a null distribution. We calculated a permuted *P*-value (*P*_perm._) defined as the number of sets with a proportion of null *trans*-eVariants proximal to at least one expressed TF/co-TF equal to or greater than the observed proportion divided by 10,000. Across distance windows from ± 70kb – 1Mb, we noted that our observed proportion was significantly higher (*P*_perm._ < 0.05) than that expected by chance. (**Fig. 5d**, **Supplementary Table 25**).

### Transcriptome wide association analyses highlight genes associated with bTB susceptibility

To assess if expression patterns in the three groups of animals (bTB+, bTB−, and AAG) were correlated to bTB susceptibility, we used MOSTWAS^58^ to generate predictive models of expression and combined these, using a TWAS approach, with SNP summary statistics from multiple GWAS data sets for bTB susceptibility in four breed cohorts (Holstein-Friesian – HF, Charolais – CH, Limousin – LM, and a multi-breed panel – MB)^33^. The SNPs in these GWAS data sets were originally mapped to the UMD3.1 genome assembly and were therefore remapped to the ARS-UCD1.2 assembly for this TWAS. We first computed 29,905, 91,822, and 1,046,632 significant (FDR *P*_adj._ < 0.01) correlations between expressed cis-eTFs/coTFs and *cis*-eGenes in the bTB+, bTB−, and AAG cohorts, respectively. We then used the *MeTWAS* function in MOSTWAS to build predictive models of expression for *cis*-eGenes within each group. In total, we trained 1,604, 2,502, and 3,957 expression models in the bTB+, bTB−, and AAG cohorts, respectively (**Table 3**). The expression patterns of these genes were significantly heritable (*P* < 0.05) and achieved a McNemar’s five-fold cross-validated predicted *R*^2^ value ≥ 0.01 within the *MeTWAS* function. For each reference group and each GWAS cohort, we conducted a weighted burden test using the MOSTWAS *BurdenTest* function to identify genes with expression patterns correlated to bTB susceptibility. For genes that were significant at a Bonferroni-adjusted *P*-value < 0.05, we conducted a permutation test conditioning on the GWAS effect size and genes with a permuted *P*-value < 0.05 were considered significantly associated with bTB susceptibility.

**Table 3:** Total number of significantly heritable (*P* < 0.05) predictive expression models (*R*^2^ > 0.01) generated for the reactor (bTB+), control (bTB−) and combined all animals (AAG) cohorts with the corresponding Bonferroni adjusted *P*-value cut-off for association and number of significant genes identified across all four GWAS data sets. Numbers in brackets indicate the number of TWAS genes significant after permutation testing.

| Group | Expression models | $P$-value cut-off | Limousin (LM) TWAS genes | Holstein-Friesian (HF) TWAS genes | Charolais (CH) TWAS genes | Multi-breed (MB) TWAS genes |
| --- | --- | --- | --- | --- | --- | --- |
| bTB+ | 1,604 | $3.12 \times 10^{-5}$ | 11 (3) | 14 (7) | 14 (9) | 27 (12) |
| bTB– | 2,502 | $2.00 \times 10^{-5}$ | 10 (3) | 30 (15) | 11 (6) | 18 (9) |
| AAG | 3,957 | $1.26 \times 10^{-5}$ | 22 (12) | 46 (31) | 24 (10) | 53 (19) |

The number of genes that were significant after Bonferroni correction, and that remained significant after the permutation procedure in each of the 12 TWAS groups are shown in **Table 3**. In total, across all four GWAS cohorts (HF, CH, LM, and MB) we identified 31, 33 and 72 TWAS genes significantly associated with bTB susceptibility in the bTB+, bTB−, and AAG cohorts, respectively (**Fig. 6**). Among the cohorts, there was little overlap between TWAS genes, with many genes emerging as breed- and expression model-specific (**Supplementary Fig. 12**). Overall, we identified 136 genes dispersed across the genome with expression patterns correlated with bTB susceptibility (**Fig. 6**). Our TWAS analysis highlighted immunobiologically relevant genes such as *RGS10, GBP4, TREML2*, and *RELT* and the full results of all TWAS associations for each reference panel are provided in **Supplementary Table 26-28**.

**Fig. 6:**
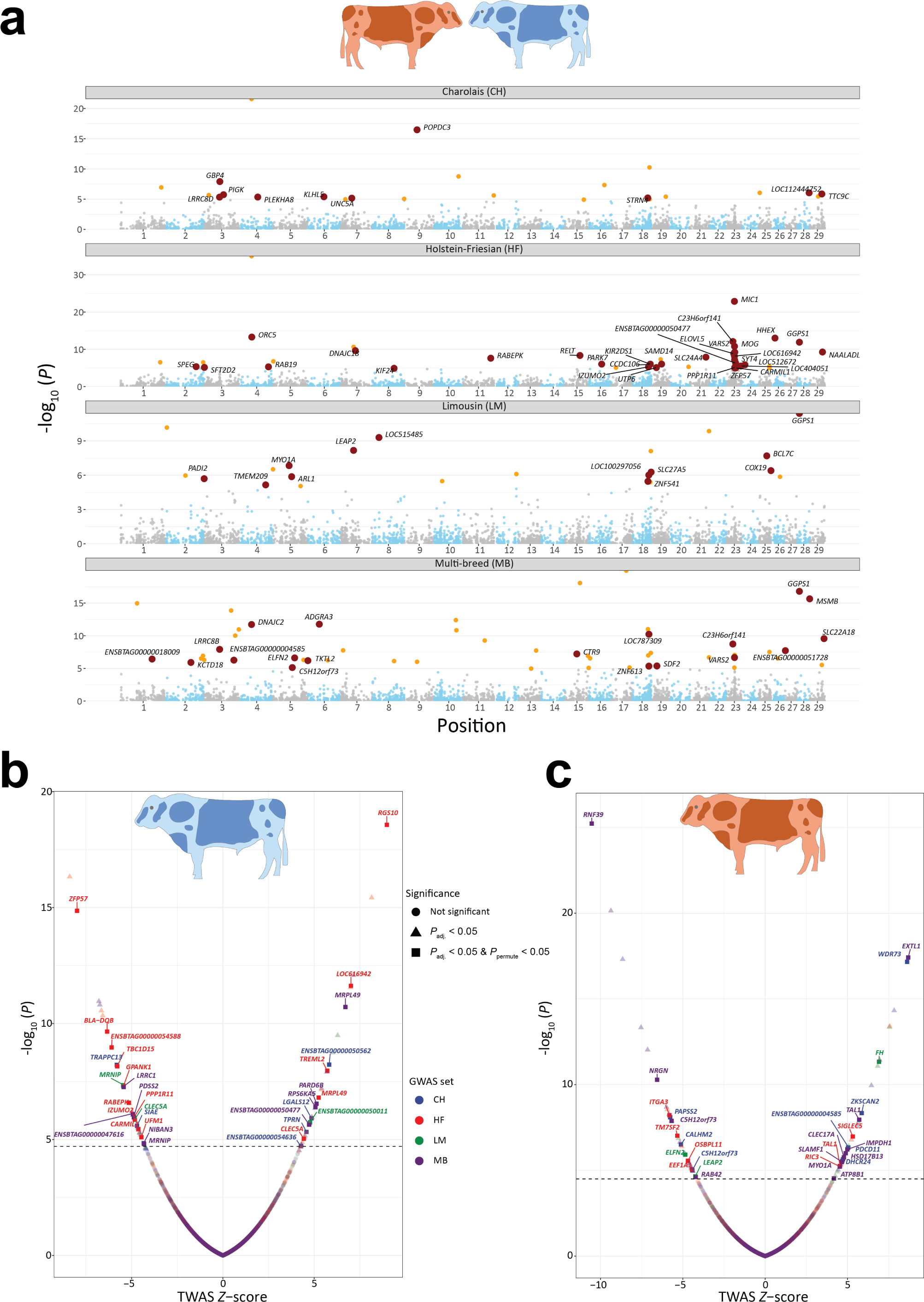
Transcriptome wide association analysis (TWAS) results. **a** Manhattan plots showing all TWAS associations for expression models generated in the analysis of all animals combined and imputed into four GWAS data sets (Charolais (CH), Holstein-Friesian (HF), Limousin (LM), and Multi-Breed (MB)). Yellow data points have a Bonferroni FDR *P*_adj._ < 0.05, and red points correspond to genes that have a Bonferroni *P*_adj._ < 0.05, and *P*_perm._ < 0.05. Labelled genes correspond to red data points in the plot. **b** Volcano plot highlighting significant TWAS associations for expression models generated in the reactor group (bTB+). The *x*-axis indicates the TWAS *Z*-score, and the *y*-axis shows the nominal (-log_10_ scale) *P*-value of association. Associations are coloured based on the GWAS data set for which the expression model was imputed into. Associations are shaped according to whether they had a Bonferroni *P*_adj._ > 0.05 (circle), *P*_adj._ < 0.05 (triangle), or *P*_adj._ ≤ 0.05 and *P*_perm._ < 0.05 (square). The dashed line corresponds to a Bonferroni *P*_adj._ cut-off (*P* < 3.12 × 10^-5^). **c** Volcano plot highlighting significant TWAS associations for expression models generated in the control group. The *x*-axis indicates the TWAS *Z*-score, and the *y*-axis shows the nominal *P*-value of association. Associations are coloured based on the GWAS data set for which the expression model was imputed into. Associations are shaped according to whether they had an FDR *P*_adj._ > 0.05 (circle), *P*_adj._ < 0.05 (triangle), or *P*_adj._ ≤ 0.05 and *P*_perm._ < 0.05 (square). The dashed line corresponds to a Bonferroni *P*_adj._ cut-off (*P* < 2.00 × 10^-5^). The figure legend for panel **b** and panel **c** is common to both.

## Discussion

We present a comprehensive multi-omics analysis, which integrates genomics, bovine PB transcriptomics and GWAS data sets for bTB susceptibility to improve our understanding of how genetic factors contribute to the interindividual variability in response to *M. bovis* infection and mycobacterial infections more broadly in a One Health context. Moreover, this study is the first application of the TWAS approach to dissecting the genomic architecture of a susceptibility trait for a mycobacterial infection that causes TB in mammals.

Bovine TB disease susceptibility is a moderately heritable quantitative trait (estimated *h*^2^ ranges between 0.08 and 0.14) with a highly polygenic and breed-specific genetic architecture that poses significant challenges for functional assignment of QTLs identified from GWAS experiments^33,76–78^. However, understanding the biology of these QTLs will be important in bridging the genome to phenome gap for bTB disease resilience because regulatory QTLs, especially *cis*- and *trans*-eQTLs, contribute a large proportion of the variance in complex trait heritability^44,45^. Additionally, it has been estimated that up to 50% of GWAS signals are shared with at least one molecular phenotype in humans^79^, with a particular enrichment observed for regulatory QTLs associated with proximal and distal gene expression regulation in PB^80^.

Analysis of differential gene expression using RNA-seq showed that the bovine PB transcriptome is substantially perturbed by *M. bovis* infection with 2,592 genes significantly (FDR *P*_adj._ < 0.05) DE (**Fig. 2c**, **Supplementary Table 5**). We detected fewer DEGs in comparison to those reported by McLoughlin, et al. ^64^ who analysed PB leukocytes from cattle infected with *M. bovis*. However, we identified more DEGs than McLoughlin, et al. ^68^ who analysed whole blood RNA-seq data from calves infected with *M. bovis* across an experimental time-course. The variability observed in this study among animals, characterised by differences in breed composition, age, duration since *M. bovis* infection, and the varied biological tissue analysed, along with the diverse cell composition associated with PB may explain the heterogenous nature of the bovine transcriptomic profile. Contrary to this, the experiments conducted by McLoughlin, et al. ^64^ and McLoughlin, et al. ^68^ featured a more controlled setting, involving Holstein-Friesian calves matched for age and breed. These observations are supported by other studies showing that population genetic structure impacts gene expression due to allele frequency differences at *cis*-eQTL sites^81^, and that ancestry effects impact the human response to viral infection in a cell type-specific manner^82^. Many of the DEGs detected here (38%) were also observed to be DE at 48 hours post infection (hpi) in bovine alveolar macrophages (bAM) challenged with *M. bovis* and were components of gene modules key to the innate immune response^83^. These shared genes included, but were not limited to, *MX2, MX1, OAS2, ISG15*, and *IRF7* that collectively constitute interferon-stimulated genes^84^. This is also reflected in our gene set enrichment analysis of highly significant DEGs where many of the top significant overrepresented functional entities were biological pathways and GO terms related to interferon signalling and induction of interferon genes (**Fig. 2d**, **Supplementary Table 6**).

Our *cis*-eQTL analysis highlighted hundreds of thousands *cis-*eVariants that were associated with thousands of *cis*-eGenes (**Table 1**), and our power of detection was dependent on sample size, which has been previously reported^53^. We also showed that there are multiple independent *cis*-eQTLs acting on thousands of genes (**Table 1**, **Fig. 3b**). Although PB is cellularly heterogeneous, we obtained good replication of *cis*-eQTLs in an external cohort from the Cattle GTEx Consortium^53^ using AC, Storey’s π_1_ statistic, and Spearman correlation of effect size estimates (**Fig. 3e**, **Fig. 3f**). While nearly all expressed genes appear to have a *cis*-eQTL in a relevant context/tissue^85^, we demonstrated that PB DEGs, which differentiate bTB+ and bTB− cattle have genomic variants associated with transcript abundance and that perturbation of these genes significantly impacts host immunobiology, most notably functions associated with defence response to virus and HIV-1 viral life cycle (**Fig. 4d**). Peripheral blood DEGs have recently been characterised as reflecting disease-induced expression perturbations rather than mechanistic disease causing changes^86^; however, the DE *cis*-eGenes identified in our study should be prioritised for further downstream functional analysis and the eVariants associated with these genes may be incorporated as prior information in future genome-enabled breeding programmes for bTB disease susceptibility traits^87,88^.

*Cis*-eQTLs explain a small proportion of expression heritability whereas *trans*-eQTLs have been estimated to contribute up to 70% of the interindividual variance in gene expression^57,89^ and tag important genomic regulatory elements and transcriptional regulators (e.g., TFs/coTFs), which will be important for bridging the genome to phenome gap in livestock species^90^. We mapped *trans*-eVariants located more than 5 Mb from the associated gene and observed an inflated number of *trans*-eGenes despite a limited sample size (*n* = 123). In humans, with a sample size of approximately 120 subjects, we would expect to detect less than five *trans*-eGenes^74^. Conversely, in the present study using 123 bovine PB transcriptomes, we detected 107 *trans*-eGenes that were associated with 5,314 *trans*-eVariants. An inflated number of *trans*-eGenes was also previously reported by the Cattle GTEx Consortium^53^ and we posit that many of these *trans*-eVariants are false positives owing to a complex intra- and interchromosomal LD relationship existing between *trans*-eVariants and top *cis*-eQTLs of the same gene. We empirically showed via permutation analysis that the LD relationship between intra- and interchromosomal *trans*-eVariants and top *cis*-eQTLs of the same gene was significantly higher than that expected by chance (**Fig. 5b**, **Fig. 5c**, **Supplementary Fig. 11c**). This LD pattern is likely a consequence of intense human-mediated selection for production traits (e.g., milk yield)^91^ and the relatively small and decreasing effective population size (*N*_e_) of European *B. taurus* breeds^92^, particularly the Holstein-Friesian breed^93^. Diminishing *N*_e_ accentuates the contribution of random genetic drift to allelic frequency changes, which can lead to random associations between loci on different chromosomes that arise at a rate inversely proportional to the *N*_e_^94^. Another evolutionary force, admixture, particularly admixture between genetically isolated populations can contribute to LD arising between unlinked sites, however such LD is likely to degrade rapidly^95–97^.

Our TWAS analyses revealed a total of 136 genes associated with susceptibility to bTB disease in cattle, many of which were breed- and expression model cohort-specific, an observation that reflects the polygenicity of this phenotypic trait^33^ (**Supplementary Fig. 12**). However, several of the genes we identified have documented roles in the host response to mycobacterial infection and the immunopathology of TB disease. For example, in our AAG-CH TWAS cohort, we identified the guanylate binding protein 4 gene (*GBP4*; *P =* 2.5 × 10^−6^; *Z* = −4.7) as being significantly associated with bTB disease susceptibility. A negative *Z*-score can be interpreted as decreased expression of this gene being associated with the trait of interest^47^. *GBP4* is an interferon-inducible gene that is upregulated and contributes to the Type 1 immune response during *M. tuberculosis* infection^98^. Additionally, expression of *GBP4* was shown to be significantly upregulated at +1 week, + 2 weeks and +10 weeks in blood samples from cattle experimentally infected with *M. bovis* and stimulated with PPD-b compared to control non-stimulated blood samples^70^ and in PB leukocytes of *M. bovis*-infected cattle compared to non-infected control animals^64^. The most significant gene associated with bTB disease susceptibility in our bTB− group was the regulator of G-protein signalling 10 gene (*RGS10*; *P* = 7.34 × 10^−18^; *Z* = 8.6), which was identified in the Holstein-Friesian GWAS panel. (**Fig. 6b**). *RGS10* encodes an important anti-inflammatory protein and has previously been implicated *in vitro* in regulating macrophage activity, specifically limiting activation of the NF-ĸB pathway, reducing expression of tumour necrosis factor (TNF), and regulating macrophage M1 to M2 repolarisation^99^. In murine models challenged with a lethal dose of influenza A virus, loss of *RGS10* resulted in increased cytokine and chemokine activity, and a more pronounced recruitment of neutrophils and monocytes to the site of infection^100^.

Members of the MTBC, including *M. tuberculosis* and *M. bovis*, have evolved a diverse range of strategies to modulate, subvert, and evade the host innate immune response and an important facet of this is manipulation of granuloma formation and function^101^. Recent multi-modal profiling of the granuloma in cynomolgus macaques (*Macaca fascicularis*) experimentally infected with *M. tuberculosis* has shown that high-burden granulomas are characterised by Type 2 immunity and tissue-protective responses that maintain essential tissue functionality while paradoxically creating a niche for bacterial persistence, whereas low *M. tuberculosis* burden granulomas are governed by an adaptive Type 1–Type 17 (Th1-Th17) and cytotoxic T cell responses that kills and destroys invading bacilli^102^. In this regard, we also identified the RELT TNF receptor gene (*RELT*; *P* = 4.4 × 10^−9^; *Z* = 5.9) in our AAG-HF TWAS cohort as being significantly associated with bTB disease susceptibility. *RELT* is a member of the TNF superfamily and evidence suggests that RELT may promote an immunosuppressive environment through suppression of IFN-γ, TNF, and IL-5 production in CD4^+^ and CD8^+^ T cells^103^. The triggering receptor expressed on myeloid like cells 2 gene (*TREML2*) was also significantly associated with bTB susceptibility in our bTB−/HF TWAS cohort (*P* = 1.1 × 10^-8^; *Z* = 5.7). In monocytes infected with *M. tuberculosis*, overexpression of *TREML2* was shown to promote *IL6* transcription through activation of STAT3 and to supress the Th1 response^104^. Expression of IL-6 induced by *M. tuberculosis* infection was also shown to inhibit the macrophage response to IFN-γ^105^ and impaired intracellular killing of mycobacteria^106^.

Taken together, based on our TWAS results, we can therefore hypothesise that the combined increased and decreased expression of several immunoregulatory genes dampens the proinflammatory immune response during *M. bovis* infection, supresses the Th1 T-cell response and contributes to macrophage M2 polarisation and Type 2 immunity characteristics that lead to bTB disease susceptibility, bacterial persistence, pathogenesis, and disease.

Although the TWAS approach is being increasingly applied to complex traits in plant and animal species, the statistical framework underpinning the methodology has come under criticism due to an inflated type 1 error rate as a consequence of failing to account for predictive expression model uncertainty in the gene-trait association test^51^. Stringent gene filtering through application of a two-step statistical significance process with a Bonferroni *P*_adj._ < 0.05 threshold followed by a post-hoc permutation test (*P* < 0.05) appears to control for this inflated false positive rate^51^. However, the permutation scheme itself is highly conservative and as such, true causal genes associated with the trait of interest may be filtered out owing to insufficient power^47^. Immunologically relevant genes that did not achieve a *P*_perm._ < 0.05, but that may be associated with bTB disease susceptibility, included the polymeric immunoglobulin receptor gene (*PIGR*) in the AAG-MB TWAS cohort (*P* = 2.83 × 10^−7^; *Z* = −5.1). *PIGR* encodes an important transmembrane receptor involved in the transport of dimeric immunoglobulin A (IgA) from the lamina propria across the epithelial/mucosal barrier enabling the production of secretory immunoglobulins that mediate innate host protection through specific and non-specific pathogen interactions^107^. Previous work has shown that *PIGR*^-/-^ mice are more susceptible to *M. tuberculosis* infection and have reduced IFN-γ and TNF expression and a delayed induction of mycobacteria-induced immune responses^108^. PTEN induced kinase 1 gene (*PINK1*) was also identified as being associated with bTB susceptibility in the bTB−/HF TWAS cohort (*P* = 7.0 × 10^−8^; *Z* = 5.4). In bovine monocyte-derived macrophages (MDM) challenged with *M. bovis*, expression of *PINK1* was shown to benefit the pathogen in the host cell through induction of mitophagy, promoting its intracellular survival by inhibiting xenophagy^109^.

While the TWAS reference panels (bTB+, bTB−, and AAG) used in this study are primarily composed of crossbred cattle, with Holstein representing the bulk of animal ancestry (**Fig. 2a**, **Fig. 2b)**, three of the GWAS data sets that were used to impute the expression models were generated from single breed population samples (Charolais, Holstein-Friesian, and Limousin). The genetic heterogeneity across the bTB+/bTB−/AAG reference panels and GWAS cohorts, therefore, makes it challenging to impute reference expression models. A reference panel that better matches the GWAS cohort would result in more power to detect genes associated with *M. bovis* infection susceptibility/resistance traits. This issue may account for the detection of more significant TWAS genes in the Holstein-Friesian GWAS cohort versus the other breed cohorts across the bTB− and AAG reference panels as Holstein was the predominant ancestry (**Table 3**). This observation aligns with results from a previous integrative genomics study. Compared to the Charolais and Limousin GWAS data sets, substantially more significant bTB susceptibility associated SNPs were detected in the Holstein-Friesian GWAS data set following integration of functional genomics outputs from Holstein-Friesian bovine AMs challenged *in vitro* with *M. bovis*^83^.

The animals in the bTB+ reference panel have a confirmed bTB diagnosis and are maintained for bTB diagnostics potency testing; therefore, it is possible that some of the results from the TWAS may be confounded by horizontal pleiotropy owing to the same causal variant having independent effects on both expression and the trait^47^. We would therefore prioritise significant TWAS associations identified in the analysis of the bTB− group for further downstream analyses. Lastly, it is challenging to evaluate causality from our TWAS results due to issues associated with sharing of GWAS variants between expression models, coregulation of a putatively causal and non-causal gene(s), and correlation of predicted expression models between tested genes^48^. A combination of a larger tissue/cell specific reference panel in conjunction with other integrative and functional genomic techniques such as colocalization^110^ and Mendelian randomisation^111^ would facilitate this approach.

## Methods

### Animal recruitment, sampling, and data acquisition

A total of *n* = 60 cattle infected with *M. bovis* and *n* = 63 non-infected control animals were recruited for the purpose of this study. All animals were male and were born between 2014 and 2019. The *M. bovis*-infected cattle (bTB+) were selected from a panel of naturally infected animals maintained for on-going tuberculosis surveillance at the Department of Agriculture, Food, and the Marine (DAFM) Backweston Laboratory Campus farm (Celbridge, Co. Kildare, Ireland). These animals were skin tested by an experienced veterinary practitioner and had positive single intradermal comparative tuberculin test (SICTT) results where the skin-fold thickness response to purified protein derivative (PPD)-bovine (PPDb) exceeded that of PPD-avian (PPDa) by at least 12 mm. As an ancillary diagnostic test carried out in series, all animals were tested for *M. bovis* infection using the whole blood IFN-γ release assay (IGRA) (BoviGAM^®^ – Prionics AG, Switzerland)^71^. During *post-mortem* examination, all the animals disclosed multiple lesions consistent with bovine tuberculosis. The non-infected control animals (bTB−) were selected from bTB-free cattle herds (all SICTT negative) and with no recent history of *M. bovis* infection.

Peripheral blood was sampled from each animal using blood collection tubes with blood harvested from the tail vein. All tubes were inverted 4-6 times immediately after sampling and transported to the laboratory in a refrigerated cool box within three hours of collection. Whole blood was collected in Tempus^™^ blood RNA stabilisation tubes (Thermo Fisher Scientific) for isolation of total RNA. All tubes intended for RNA extraction were stored at −80°C prior to isolation and purification. A single heparin-coated tube was also collected from each animal (bTB+ and bTB−) for same-day IGRA testing to confirm infection status at the UCD Tuberculosis Diagnostics and Immunology Research Centre. Further details on genomic and transcriptomic data acquisition are detailed in **Supplementary Note 1**.

### Basic population genomics analysis and imputation

All animals in the study were genotyped using the Affymetrix Axiom^™^ Genome-Wide BOS-1 Array (Thermo Fisher Scientific) with SNP positions originally mapped to the UMD3.1 bovine reference genome assembly^112^. The CEL files were imported into Axiom Analysis Suite software tool v.5.1.1.1 following the Axiom Best Practices Genotyping Analysis Workflow with the required sample attributes^113^. The SNPolisher Recommended Probesets were exported in PLINK format annotated with the Axiom_GW_Bos_SNP_1.na35.annot.db annotation database using genome version UMD 3.1 and NCBI version 6. This analysis yielded a total of 591,947 SNPs (91.23%) for downstream analyses. Prior to remapping of SNPs to the current ARS-UCD1.2 genome build^114^, the genetic structure and diversity of the study population was evaluated as follows. PLINK v1.90b6.25^115^ was used to filter out SNPs with a minor allele frequency (MAF) < 10%; that deviated from Hardy-Weinberg equilibrium (HWE; *P* < 1 × 10^-6^); and with a call rate < 0.95. The PLINK -- *indep-pairwise* command was then used to prune variants in linkage disequilibrium (LD) with the following parameters: window size = 1000 kb; step size = 5 variants; and *r*^2^ > 0.2. Following these steps there were 34,272 SNPs available for examination of genetic structure using ADMIXTURE v.1.3^116^ and principal component analysis (PCA) using PLINK.

The ADMIXTURE analysis was performed with the *--cv* option such that setting the number of ancestral populations to *K* = 2 produced the lowest cross-validation error. We then used pophelper v.2.3.1^117^ to generate a structure plot. For the PCA analysis, we used the --*pca* function in PLINK and used a custom R v.4.3.2^118^ script to plot the PCA results using ggplot2 v.3.4.1^119^.

For the imputation up to whole-genome sequence (WGS) scale data, raw genotyped variants were first remapped from UMD3.1 to the ARS-UCD1.2 bovine genome assembly (**Supplementary Note 2**). Following this, a global cattle reference panel from Dutta, et al. ^72^ was used as the imputation reference panel, which comprised a total of 10,282,187 SNPs derived from *n =* 287 distinct animals spanning a diverse range of breeds and geographic locations (55 populations: 13 European, 12 African, 28 Asian, and two Middle Eastern). Imputation was performed using Minimac4 v.1.03^120^ with default parameters to impute the target genotype data set up to WGS, which resulted in a master imputed data set consisting of all *n =* 123 animals with genotypes for 10,282,037 SNPs (**Supplementary Note 2**).

### Transcriptomics data quality control, read alignment, and read mapping

The paired-end RNA-seq FASTQ files (*n* = 123; 60 bTB+ and 63 bTB−) was assessed using FastQC v.0.11.5^121^, which showed that the RNA-seq data set was of sufficiently high quality to negate the requirement for hard or soft trimming. Following this, RNA-seq reads were aligned to the ARS-UCD1.2 bovine reference genome using STAR v.2.7.1a^122^. Read counts for each gene were then quantified using featureCounts v.2.0.6^123^ and the ARS-UCD1.2 ensemble annotation file (https://ftp.ensembl.org/pub/release-110/gtf/bos_taurus/Bos_taurus.ARS-UCD1.2.110.gtf.gz) excluding chimeric fragments, aligning reads in a reversely stranded manner, and considering only fragments with both ends successfully aligned for quantification.

### Missing data imputation and sample mismatch assessment

Control sample C028 did not have any date of birth information available (**Supplementary Table 1**). Therefore, we inferred the age of C028 as the mean of all other animals that were sampled on the same date (02/05/2017). Control samples C039 and C041 were assigned the same animal identification number; therefore, to ensure that these animals were not duplicates, we estimated the identity-by-state (IBS) distance values between all samples by using the pruned SNPs prior to imputation to identify and remove duplicate animals using PLINK. The IBS distance values were calculated as:

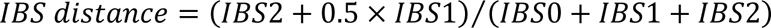

where *IBS0* is the number of IBS 0 non-missing variants, *IBS1* is the number of IBS 1 non-missing variants and *IBS2* is the number of IBS 2 non-missing variants. Sample pairs with IBS distance values > 0.85 were considered duplicates and only one sample was retained for subsequent analyses^53^.

To ensure that the transcriptomics data and genome-wide SNP data for all 123 animals (bTB− and bTB+) were matched, we assessed the genotype consistency using the match BAM to VCF (MBV) function^124^ that is part of the QTLtools (v 1.3.1) package^125^. Briefly, MBV reports the proportion of heterozygous and homozygous genotypes (for each sample in a VCF file) for which both alleles are captured by the sequencing reads in all BAM files. Correct sample matches can then be verified, as they should have a high proportion of concordant heterozygous and homozygous sites between the genotype data and the mapped sequencing reads.

### Differential expression analysis

A differential expression analysis (DEA) was conducted between the control (bTB−) and reactor (bTB+) animal groups using DESeq2 v.1.40.2^126^ and a design matrix, which included the following covariates: age in months, RNA-seq sequencing batch, and genetic structure in the form of PC1 and PC2 from the PCA of the pruned SNP data set prior to imputation with reactor status as the variable of interest. The PC1 and PC2 covariates were included because the crossbred/multibreed nature of the animals in our study population should be incorporated in the DEA contrast for the bTB− and bTB+ animal groups. Genes with raw expression counts ≥ 6 in at least 20% of samples were retained prior to the DEA. For the DEA, the null hypothesis was that the logarithmic fold change (LFC) between the control and the reactor group, for the expression of a particular gene is exactly 0. To account for potential heteroscedasticity of LFCs, we implemented the approximate posterior estimation for generalised linear model coefficients (APEGLM) method^127^ using the *lfcShrink* function. Genes with a Benjamini-Hochberg (BH) false discovery rate (FDR) adjusted *P*-value^128^ (*P*_adj._) < 0.05 and a LFC > 0 or < 0 were considered significantly differently expressed (DE).

### *Cis*-eQTL mapping

For the mapping of *cis*-eQTLs, we used the human GTEx Consortium^74^ pipeline with some minor modifications. We conducted the *cis*-eQTL analysis on the control group (bTB−), the reactor group (bTB+), and a combined group of all 123 animals (AAG). Raw RNA-seq read counts were normalised using the trimmed mean of the M values (TMM) method^129^ and the expression values for each gene were then inverse normally transformed across samples to ensure the molecular phenotypes followed a normal distribution. Genes with raw expression counts ≥ 6 and a transcript per million (TPM)^130,131^ normalised expression count ≥ 0.1 in at least 20% of samples were retained for the eQTL analysis. For each group, we used the PCAForQTL R package v.0.1.0^132^ to identify hidden confounders in the normalised and filtered expression matrices. The number of latent variables selected was determined using the elbow method via the *runElbow* function in PCAForQTL. We then merged these inferred covariates with known covariates (the top five genotype PCs of the imputed data set, age in months, sequencing batch, and infection status, where applicable) and removed highly correlated known covariates captured well by the inferred covariates (unadjusted *R*^2^ ≥ 0.9) using the PCAForQTL *filterKnownCovariates* function.

For the *cis*-eQTL mapping procedure, we used TensorQTL v.1.0.8^133^. We defined the *cis* window as +/− 1 Mb from the transcriptional start site (TSS) of a gene. To identify significant *cis*-eQTLs, we invoked the permutation strategy in TensorQTL^134^ to estimate variant-phenotype associated empirical *P-*values with the parameter --mode *cis* to account for multiple variants being tested per molecular phenotype. We then used the Storey and Tibshirani FDR procedure^135^ to correct the *beta* distribution-extrapolated empirical *P-*values to account for multiple phenotypes being tested genome-wide. A gene with at least one significant associated *cis*-eQTL was considered a *cis*-eGene.

To identify significant *cis*-eVariants associated with detected *cis*-eGenes, we followed the procedure implemented by the Pig GTEx Consortium^136^. Briefly, we first obtained nominal *P*-values of association for each variant-gene pair using the parameter --mode *cis_nominal*. We then defined the empirical *P*-value of a gene which was closest to an FDR of 0.05 as the genome wide empirical *P-*value threshold (*pt*). Next, we calculated the gene-level threshold for each gene from the beta distribution by using the *qbeta(pt*, *beta_shape1*, *beta_shape2*) command in R with beta_shape1 and beta_shape2 being derived from TensorQTL. Variants with a nominal *P*-value of association below the gene-level threshold were included in the final list of variant-gene pairs and were considered as significant *cis*-eVariants.

Following the Pig GTEx Consortium^136^, to identify genes with multiple independent-acting *cis*-eQTLs, we performed a conditional stepwise regression analysis using the parameter --mode *cis_independent*. Briefly, the most significant variant was considered a putative *cis*-eQTL if it had a nominal *P*-value below the genome-wide FDR threshold inferred above. Next, using a forward stepwise regression procedure, the genotypes of this variant were residualized out from the phenotype quantifications and the process of regression, selection, and residualization was repeated until no more variants were below the *P*-value threshold resulting in *n* independent signals per gene. Finally, using backward stepwise regression, nearby significant variants were assigned to inferred independent signals.

### *Trans*-eVariant mapping and permutation analysis

We conducted *trans*-eQTL mapping on all three groups of animals using QTLtools v.1.3.1^125^ using the --*trans* option. We first tested all variant phenotype pairs using the *--nominal* and *--normal* parameters together, including the same covariates described for the *cis*-eQTL mapping procedure and reported those with a nominal association below a threshold of *P* < 1 × 10^-5^ and that were not proximal (< 5 Mb) to the tested phenotype. We then characterised the null distribution of associations by employing the --*permute* option and used the QTLtools *runFDR_ftrans.R* script to estimate the FDR. Briefly, the nominal and permuted *P*-values are ranked in descending order and the FDR for a particular variant-phenotype pair is calculated by counting the number of permutation hits with smaller *P*-values than the nominal *P*-value for a variant-phenotype pair and, finally, dividing this number by the rank of the pair. Variants with an FDR < 0.05 were considered significant *trans*-eVariants. Given the small number of *trans*-eGenes identified in the control (bTB−) and reactor (bTB+) cohorts, we decided to focus on the larger cohort (bTB− and bTB+) for analysis of *trans*-eVariants.

We hypothesized that top intra and interchromosomal *trans*-eQTLs were in high LD with top *cis*-eQTLs of the same gene. To test this hypothesis, we performed a permutation analysis where we randomly sampled 10,000 sets of null intrachromosomal variant pairs and interchromsoomal *trans*-eVariants respectively. For the intrachromosomal set, we computed the LD (*r*) between each set and compared the distribution of the means and medians to our observed distribution. For the interchromsomal set, we computed the LD (*r*) between null intrachromsomal *trans*-eVariants and top *cis*-eQTLs of the same gene and compared the means and medians of the 10,000 sets to our observed distribution. For both the inter and intrachromosomal LD analyses, we calculated a permuted *P*-value (*P*_perm._) defined as the number of sets with a mean or median LD (*r*) value respectively greater than or equal to our observed LD values divided by 10,000. A more detailed description of this analysis is outlined in **Supplementary Note 3**.

Lastly, we hypothesised that the remaining top *trans*-eVariants were proximal to expressed transcription factors (TFs) or transcription factor co-factors (co-TFs). To empirically test this, we first removed *trans*-eVariant gene pairs if the top *trans*-eVariant was in LD (*r*^2^ > 0.01) with the top *cis*-eVariant associated to the same gene and only considered remaining *trans*-eVariants that were highly significant (FDR < 0.01). We then downloaded genomic coordinates for annotated TFs/co-TFs from the AnimalTFDB: v.4.0 database^75^. We calculated the proportion of the filtered top *trans*-eVariants that were proximal to at least one expressed TF/co-TF at genomic intervals ranging from ±10 kb to ±1 Mb versus 10,000 random sets of SNPs to generate a null distribution. For each distance window, we obtained a *P*_perm._ value defined as the number of sets with a proportion of null *trans*-eVariants proximal to at least one expressed TF/co-TF equal to or greater than the observed proportion divided by 10,000.

### Replication of *cis-*eQTLs

To assess the replicability of *cis*-eQTLs identified for each group in an independent cohort, we first downloaded blood *cis*-eQTL summary statistics (both permuted and nominal associations) from the Cattle GTEx Consortium (https://cgtex.roslin.ed.ac.uk/wp-content/plugins/cgtex/static/rawdata/Full_summary_statisitcs_cis_eQTLs_FarmGTEx_cattle_V0.tar.gz). We used three different measurements of agreement of eQTL effects when comparing eQTLs across the two studies: allele concordance (AC), π_1_ and Spearman correlation (*ρ*). AC provides an indication of the proportion of effects that have a consistent direction of effect (slope) within the set of eQTLs that is significant in both the discovery (here, denoted as the control bTB−, reactor bTB+, and combined (AAG; bTB− and bTB+ cohorts) and the replication cohort (the Cattle GTEx) and is expected to be 50% for random eQTL effects^137^. The parameter π_1_^135^ represents the proportion of true positive eQTL *P*-values in the replication cohort and is calculated as 1 – π_0_ (the proportion of true null eQTL *P*-values). The Spearman *ρ* statistic estimates the correlation between the effect sizes (slope) of significant eQTLs in the discovery cohort and matched associations in the replication cohort, regardless of significance in the latter.

To calculate AC, we matched significant eQTLs in the discovery cohort to significant eQTLs in the replication cohort. We then calculated the proportion of these eQTLs that showed the same direction of effect. To calculate π_1_, we obtained the *P*-values in the replication cohort of significant associations identified in the discovery cohort and used the *qvalue* function in R to estimate π_0_. We then calculated π_1_ as 1 – π_0_. Uncertainty estimates of π_1_ were obtained using 100 bootstraps where SNPs were sampled with replacement and π_1_ was recomputed each time^138^. To obtain the Spearman *ρ* statistics, we calculated the Spearman correlation between significant eQTLs identified in this study to matched variant:gene pairs in the replication cohort, regardless of significance.

### GWAS data pre-processing

GWAS summary statistics for the present study were obtained from a single and multi-breed GWAS experiment that leveraged WGS data from Run 6 of the 1000 Bull Genomes Project^139^ as an imputation reference panel. The GWAS used estimated breeding values (EBVs) derived from an *M. bovis* infection phenotype as the trait of interest for *n* = 2,039 Charolais, *n* = 1,964 Limousin, and *n* = 1,502 Holstein-Friesian cattle^33^. Variants were remapped from UMD 3.1 to ARS-UCD1.2 using a custom R script that was developed for a previous study that integrated the GWAS summary statistics with functional genomics data obtained from *M. bovis*-infected bovine alveolar macrophages (bAM)^83^. To check for instances of strand flips, the reference and alternative allele pairs derived from Run 6 of the 1000 Bull Genomes Project were compared to reference and alternative allele pairs in the ARS-UCD1.2 reference genome (https://sites.ualberta.ca/~stothard/1000_bull_genomes/ARS1.2PlusY_BQSR.vcf.gz). If a strand flip occurred, the beta values for each SNP were also inverted. A Wald-statistic *Z* score for each GWAS SNP was calculated by dividing the effect size (*β*) of a SNP with the standard error of the effect size.

### Transcriptome-wide association study (TWAS) analysis

Imputed genotype data for the three groups were converted to binary (.bed) format using PLINK with the *--keep-allele-order* parameter. The resulting files were then loaded into R using the bigsnpr v.1.10.8 and bigstatsr v.1.5.6 R packages^140^. Predictive models of expression for each gene were generated using the Mediator-enriched TWAS (MeTWAS) function within the MOSTWAS package v.0.1.0^58^. Briefly, MeTWAS first identifies an association between a mediating biomarker (e.g., a TF) and a gene of interest. It then builds a predictive model of expression for the mediating biomarker considering SNPs local to the biomarker. The predicted expression pattern of the biomarker (determined via five-fold cross-validation) is then included as a fixed effect with the effect sizes of putative mediators on the expression levels of the gene of interest estimated by ordinary least squares regression. Lastly, for the final predictive model of the gene of interest, the *cis*-eVariants are fitted as random effects using either elastic net regression or linear mixed modelling, whichever produces the highest five-fold McNemar’s cross-validated adjusted *R*^2^ value.

The mediating biomarkers used in MeTWAS included expressed regulatory proteins (TFs and co-TFs) curated from the AnimalTFDB database^75^. We first computed associations between mediating biomarkers and genes through correlation analysis with significant associations (BH-FDR < 0.01) being retained. We then retained mediating biomarker:gene associations in instances where the mediating biomarker was considered a *cis*-eGene. Genes that had significant non-zero heritabilities (nominal *P <* 0.05) for their expression levels, as computed by the likelihood ratio test (LRT) from the genome-wide complex trait analysis (GCTA) software tool v.1.94.1^141^ and for which MOSTWAS-derived predictive models achieved a five-fold McNemar’s cross-validated adjusted *R*^2^ value *≥* 0.01 were retained for the gene–trait association test. The maximum number of mediating biomarkers to include in the expression model for a gene was set to ten.

Within the MOSTWAS framework, expression models were imputed into the GWAS summary statistics using the ImpG-Summary algorithm^47,142^ and a weighted burden *Z*-test was employed in the gene–trait association test^47,142^. Genes with a Bonferroni-adjusted *P*-value < 0.05 were considered candidate genes associated with bTB susceptibility. To assess whether the same distribution of GWAS SNP effect sizes could yield a significant association by chance, we implemented a permutation scheme on significant (Bonferroni-adjusted *P*-value < 0.05) TWAS genes where we sampled, without replacement, the SNP effect sizes 1000 times and recomputed the weighted burden test statistic to generate a permuted null distribution^47^. Genes with a permuted *P*-value < 0.05 were considered significantly associated with bTB disease status.

### Gene set overrepresentation and functional enrichment analyses

Gene set overrepresentation and functional enrichment analyses was conducted using a combination of the g:GOSt tool within g:Profiler v.0.2.2^143^ and Ingenuity^®^ Pathway Analysis – IPA^®^ (Summer 2023 release; Qiagen). For IPA^®^, the target species selected included *Homo sapiens*, *Mus musculus*, and *Rattus rattus* with all cell types selected in addition to the Experimentally Observed and High Predicted confidence settings. We followed best practice recommendations to account for tissue-specific sampling biases in gene set overrepresentation and functional enrichment analyses^144^. Consequently, for analysis of differentially expressed genes (DEGs), the background set consisted of all expressed genes that were tested for differential expression. For analyses of genes between the eQTL and DEGs, the background set consisted of the intersection between the genes tested in both analyses. For g:Profiler, the organism selected was *B. taurus* and an ordered query list (based on the adjusted *P*-value from the differential expression analysis) was inputted. For analyses of our query gene sets, we selected the gene ontology biological processes (GO:BP) and the cellular component (GO:CC)^145^ databases in addition to the Kyoto encyclopaedia of genes and genomes (KEGG)^146^ and Reactome^147^ repositories. To identify significantly enriched/overrepresented pathways, a BH-FDR multiple testing correction was applied (*P*_adj._ < 0.05).

### Computational infrastructure and reproducibility

All data-intensive computational procedures were performed on a 36-core/72-thread compute server (2× Intel^®^ Xeon^®^ CPU E5–2697 v4 processors, 2.30 GHz with 18 cores each), with 512 GB of RAM, 96 TB SAS storage (12 × 8 TB at 7200 rpm), 480 GB SSD storage, and with Ubuntu Linux OS (version 18.04 LTS).

## Declarations

### Ethics approval and consent to participate

All experimental procedures involving animals were conducted under ethical approval from the University College Dublin (UCD) Animal Research Ethics Committee (AREC-19-09-MacHugh) and experimental license AE18982/P141 from the Irish Health Products Regulatory Authority (HPRA) in accordance with the Cruelty to Animals Act 1876 and in agreement with the European Union (Protection of Animals Used for Scientific Purposes) regulations 2012 (S.I. No.543 of 2012).

### Consent for publication

Not applicable.

### Availability of data and materials

The RNA-seq data from the 60 *M. bovis*-infected (bTB+) and 63 control (bTB−) cattle is available at the Gene Expression Omnibus (GEO) with the BioProject Accession GSE255724. The raw UMD 3.1 high density genotype data and ARS UCD1.2 imputed and filtered data is available at 10.5281/zenodo.10658453. GWAS summary statistics data were obtained from the Irish Cattle Breeding Federation (ICBF) and additional information about sequence and genotype data availability is provided by Ring et al.^33^. The computer code and scripts used in this study are available at the following GitHub link: https://github.com/jfogrady1/BovineQTL.

### Author contributions

J.F.O.G, K.G.M, E.G, I.C.G, S.V.G, and D.E.M conceived and designed the study. D.E.M, I.C.G, S.V.G, J.G.D.P, and E.L.C acquired funding for the study. E.G, K.G.M and M.M facilitated access to animals for the study and C.N.C, G.P.M, S.L.F.O.D, J.A.W and J.A.B performed experimental work. J.F.O.G performed all bioinformatics analyses in collaboration with J.G.D.P, V.R, H.P, J.A.W, G.P.M., and T.J.H. and V.R. supplied the WGS Global Reference Panel and provided advice for variant remapping, strand flipping, and genomic imputation. H.P contributed to biological interpretation of results and discussions around *trans*-eQTL mapping. J.F.O.G wrote the first draft of the manuscript, and prepared all figures, tables, and supplementary information with input from D.E.M. All authors read and approved the final manuscript.

## Acknowledgements

The authors would like to thank Mairead Doyle for their help in interpreting the interferon-γ release assay test results for the animals used in the study. The authors would also like to express their gratitude to Bojan Stojkovic for assistance with animal ethics and Di Wang for assistance with experimental work.

