## Supplementary Information for "Integrative genomics sheds light on the immunobiology of tuberculosis in cattle"

### Supplementary Note 1

#### Genomic and transcriptomic data generation

Genomic DNA was extracted and purified for each animal (bTB+ and bTB−) using 200 µl of whole blood and the QIAamp® DNA Blood Mini Kit (Qiagen) following the manufacturer's recommendations. Purified DNA was quantified using a NanoDrop™ One microvolume spectrophotometer (Thermo Fisher Scientific) and stored at −20ºC. The extracted DNA was used for SNP genotyping using the Axiom™ Genome-Wide BOS 1 Bovine Array (Thermo Fisher Scientific), which assays 648,315 SNPs across the bovine genome. SNP array genotyping was performed by an external provider (IdentiGEN/MSD Animal Health).

Total RNA preparation was performed for each animal (bTB+ and bTB−) using the Tempus™ Spin RNA Isolation Kit (Thermo Fisher Scientific). The Tempus™ tubes were inverted to mix the contents, which were then poured into sterile 50 ml conical tubes and total RNA was isolated and purified according to the manufacturer's instructions. Following total RNA purification, an Agilent 2100 Bioanalyzer and the RNA 6000 Nano LabChip kit (Agilent Technologies, Inc.) were used to quality check and generate RNA integrity number (RIN) values for all samples, which were then stored at −80°C. Total RNA samples (bTB+ and bTB−) were used with the NEBNext® Ultra™ RNA Library Prep Kit for Illumina® (New England Biolabs) to construct sequence barcode-indexed RNA-seq libraries, which were then used for 150 bp paired-end sequencing on the NovaSeq™ 6000 Sequencing System (Illumina, Inc.). RNA-seq library preparation and sequencing were performed by an external provider (Novogene).

### Supplementary Note 2

#### Variant remapping and strand flipping

Genomic coordinates of variants were updated from UMD3.1 to the ARS-UCD1.2[1] bovine genome assembly using liftOver implemented in the R package rtracklayer v.1.54.0[2] and only SNPs were retained that matched the positions in a file of the Affymetrix Axiom™ Genome-Wide BOS-1 array coordinates updated to ARS-UCD1.2 available from the United States Department of Agriculture National Animal Genome Research Program (USDA-NAGRP) data repository (www.animalgenome.org/repository/cattle/UMC_bovine_coordinates). Due to potential strand flipping issues, all remaining palindromic SNPs were removed. Raw SNP genotype files with updated coordinates were then filtered using PLINK v1.90b6.25 to remove animals with a genotype call rate < 0.95 and to restrict analysis to autosomal SNPs with a call rate > 0.95.

Strand mismatches or miscoding of alleles can significantly reduce imputation performance[3] and statistical power in the context of genomic association studies[4]. To identify the alleles and associated genotypes that needed to be flipped, original reference and alternative allele pairs for each SNP were first determined using the Axiom™ Genome-Wide BOS-1 Array master annotation file (www.thermofisher.com/order/catalog/product/sec/assets?url=TFS-Assets/LSG/Support-Files/Axiom_GW_Bos_SNP_1-na35-annot-csv.zip) due to issues associated with the encodings of the major and minor alleles in PLINK. Following this, the reference and alternative allele pairs of the filtered VCF file were compared to the USDA-NAGRP BOS-1 ARS-UCD1.2 reference allele file (www.animalgenome.org/repository/cattle/UMC_bovine_coordinates). If the NAGRP reference allele matched the alternative allele or the complement of the alternative allele in the filtered VCF file, the SNP and associated genotypes were flipped. By following guidelines detailed in Winkler, et al. [4], the allele frequencies of SNPs calculated from all $n$ = 123 animals in the filtered genotype VCF file were compared to the allele frequencies of matched SNPs from a total of $n$ = 56 *Bos taurus* European animals. To identify SNPs with spurious allele frequency patterns, the *SmoothScatter* plotting function in R was used to identify and remove a total of $n$ = 150 SNPs that deviated from the line of parity.

### Genome-wide SNP imputation

For the imputation up to whole-genome sequence (WGS) scale data, a global cattle reference panel was used, which comprised a total of 10,282,187 SNPs derived from $n$ = 287 distinct animals spanning a diverse range of breeds and geographic locations (55 populations: 13 European, 12 African, 28 Asian, and two Middle Eastern). Details of how the original WGS data were processed is provided by Dutta, et al. [5] and the quality control procedure implemented on the data set is described by Riggio, et al. [6]. Briefly, Illumina WGS data for $n$ = 427 animals representing global cattle breeds were aligned to the UCD-ARS1.2 genome. Animals and SNPs with a high proportion of missingness (> 25%), SNPs with a poor genotype quality (GQ) (< 25) and highly related individuals (relatedness value estimated from VCFtools v.0.1.15[7] with *-relatedness2*[8] > 0.0625) were removed leaving a total of $n$ = 287 animals with 10,282,187 variants. The 150 genotyped SNPs with spurious allele frequency patterns were also removed from the WGS reference set prior to imputation.

The VCFtools package was used to split the reference and target data sets into 29 VCF files representing the 29 bovine autosomes. Beagle v.5.4[9] was then used to phase the reference and target data sets separately and Minimac3 v.2.0.1[10] was used with default settings to generate the reference haplotype m3.vcf files. Following this, Minimac4 v.1.03[10] was used with default parameters to impute the target genotype data set up to WGS scale and Bcftools v.1.7[11] was used to concatenate all 29 imputed VCF files, which resulted in a master imputed data set consisting of all $n = 123$ animals with genotypes for 10,282,037 SNPs.

### Supplementary Note 3

#### Permutation analyses of intra- and interchromosomal trans-eQTL results

We first performed an assessment of intrachromosomal *trans*-eVariants detected with our analysis of all animals. We hypothesised that a strong LD pattern was present between top intrachromosomal *trans*-eVariants and top *cis*-eQTLs of the same gene. To test this hypothesis, we first calculated the LD relationship between the top intrachromosomal *trans*-eVariants and *cis*-eQTLs of the same gene. We then randomly sampled and estimated the LD relationship for 50,000 variants to obtain a reasonable sample size. We then filtered for variants which were ≥ 5 Mb and < 2 standard deviations of the observed distribution away from each other. This distribution represented the distance between the top *trans*-eVariants and *cis*-eQTLs of the same gene. This effectively generated a null distribution of the LD relationship between intrachromosomal variants for this distance window. We then randomly sampled, with replacement, a total of 10,000 sets from this pool of variants and calculated the mean and median LD relationship between null intrachromosomal *trans*-eVariants and top *cis*-eQTLs for each of these sets. For these two distributions, we calculated a permuted *P*-value ($P_{\text{perm.}}$) defined as the number of sets with a mean or median LD ($r$) value respectively greater than or equal to our observed LD value divided by 10,000.

We next focused on interchromosomal *trans*-eVariants. Again, we hypothesised that a complex interchromosomal LD pattern existed between interchromosomal *trans*-eVariants and *cis*-eQTLs of the same gene. To test this hypothesis, we first computed the LD relationship between the top significant interchromosomal *trans*-eVariants and top *cis*-eQTL of the same gene. For each *trans*-eQTL, we randomly sampled with replacement a total of 1,000 SNPs from the same chromosome and same allele frequency as putative *trans*-eVariants and computed the LD relationship between these variants and the *cis*-eQTLs. We then employed a similar permutation scheme as outlined above where we randomly sampled 10,000 sets of *trans*-eVariants and calculated the median and mean LD ($r$) value for these sets of null interchromosomal *trans*-eVariants and *cis*-eQTLs. For these two distributions, we calculated a $P_{\text{perm.}}$ value defined as the number of sets with a mean or median LD ($r$) value respectively greater than or equal to our observed LD value divided by 10,000.

### Supplementary Note 4

#### Variant remapping and genome-wide SNP imputation analysis

For the remapping procedure of variants from the UMD 3.1 to the ARS-UCD1.2 *B. taurus* genome build, we first removed 18,792 (3.17%) palindromic SNPs from the raw SNP-array data that was comprised of 591,947 SNPs. Autosomal variants were then remapped using a combination of methods to the ARS-UCD1.2 *B. taurus* reference genome. This included lifting over variants to the new genome build, flipping genotypes and alleles where strand flips occurred and comparing allele frequencies of matched SNPs to an external cohort of European animals. A total of 553,519 genotyped autosomal SNPs (93.5% of the original set) were remapped successfully. All SNPs had a call rate > 0.95 and all animals had a genotype call rate > 0.95. Of the remapped variants, 100,888 SNPs (18.23%) were flipped due to ambiguity associated with reference and alternative allele strands. To verify that the strand flipping procedure was implemented correctly, the UCD 1.2 reference allele frequencies of the target set were compared to 56 European animals derived from a WGS Global Reference Panel[5] at matched genomic loci. A total of 236,325 SNPs overlapped between the two groups, and we observed a significant positive correlation between the allele frequencies present in both groups (Spearman correlation ($\rho$) = 0.98; $P < 2.2 \times 10^{-16}$) (**Supplementary Fig. 4**). A total of 150 SNPs present in the target and European WGS reference set were identified as potentially harbouring spurious allele frequency patterns through substantial deviation from the line of parity. These SNPs were removed from the target set and WGS Global Reference Panel. Hence, the number of SNPs remaining in the target set and the WGS Global Reference Panel prior to imputation was 553,369 and 10,282,037, respectively.

All animals in the target set were imputed up to WGS-scale using Minimac4. Imputation performance across the entire *B. taurus* genome was assessed using the squared correlation ($R^2$) metric for imputed genotyped and untyped variants and the empirical squared correlation (E$R^2$) for imputed genotyped variants only.

### Supplementary Figures

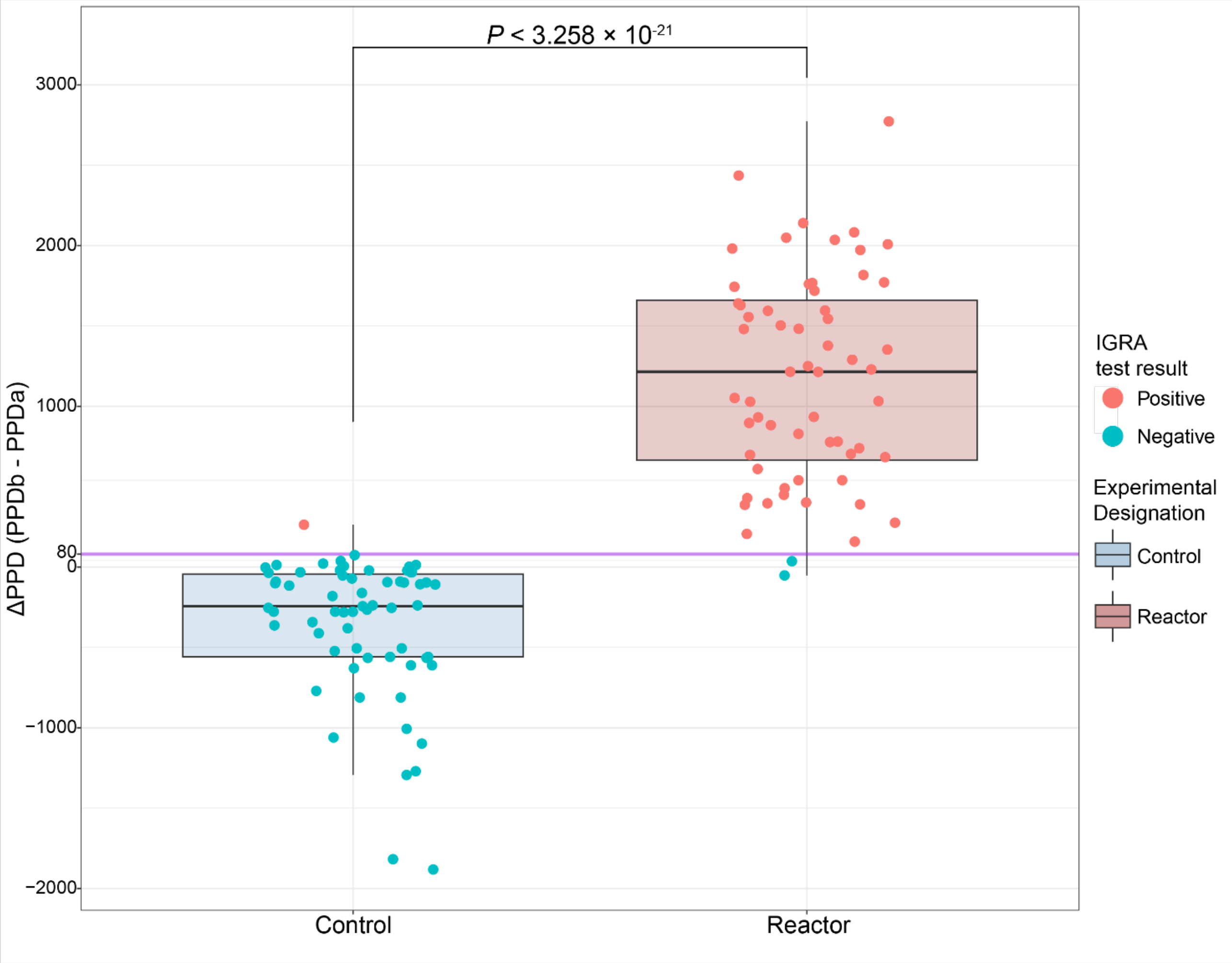

**Fig. S1: IFN-γ release assay (IGRA) test results for all (*n* = 123) animals in the present study**. Interferon-gamma (IFN-γ) release assay (IGRA) test results for animals designated as control (bTB−; *n* = 63) or reactor (bTB+; *n* = 60) animals. The *y*-axis denotes the Δ purified protein derivative (ΔPPD) value, calculated as PPD-bovine (PPDb) IFN-γ value minus the PPD-avian (PPDa) IFN-γ value. The purple line indicates the threshold for determining whether an animal is positive or negative for the IGRA test. Horizontal lines inside the boxes show the medians, Box bounds show the lower quartile (Q1, the 25th percentile) and the upper quartile (Q3, the 75th percentile).

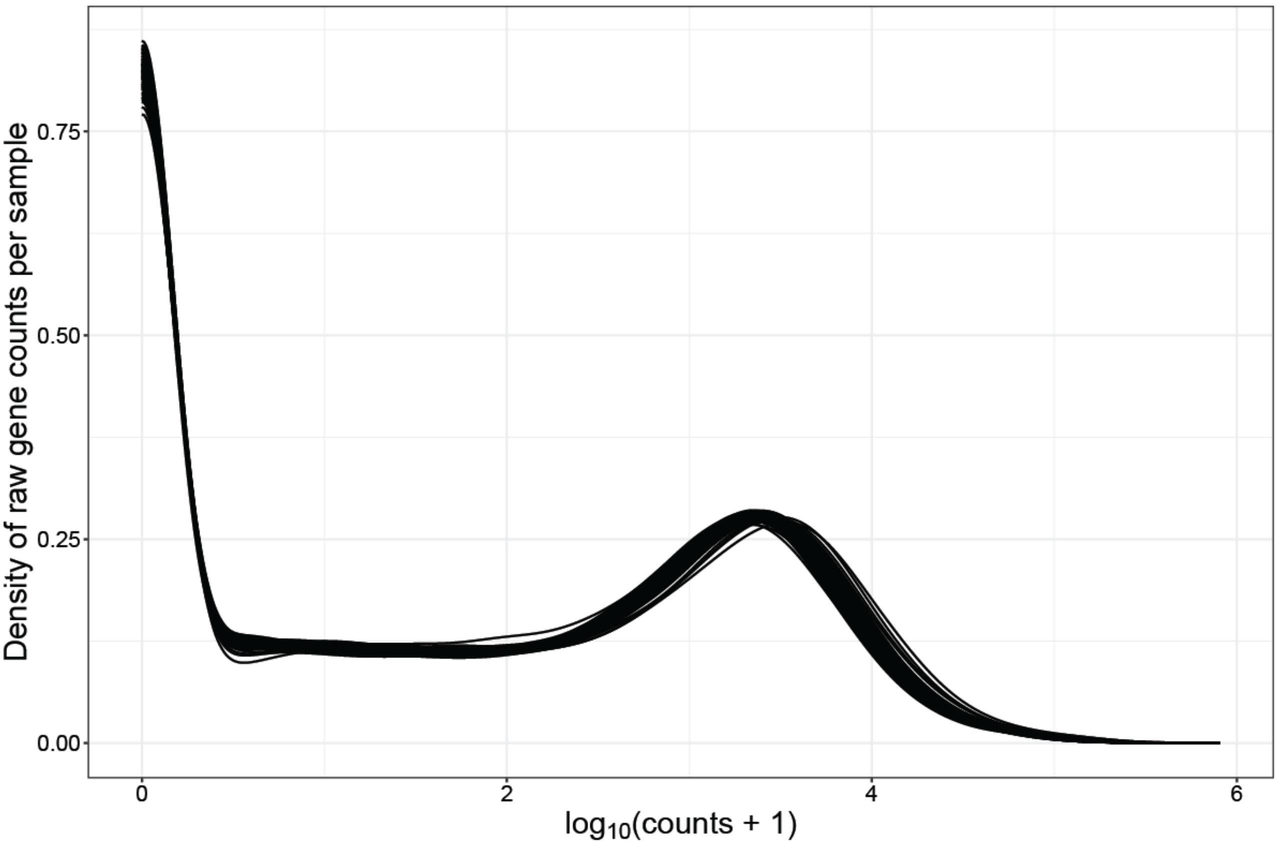

**Fig. S2: Transcriptomics data quality control.** Density of $\log_{10}$ gene counts per library ($n$ = 123) prior to filtering of lowly expressed genes.

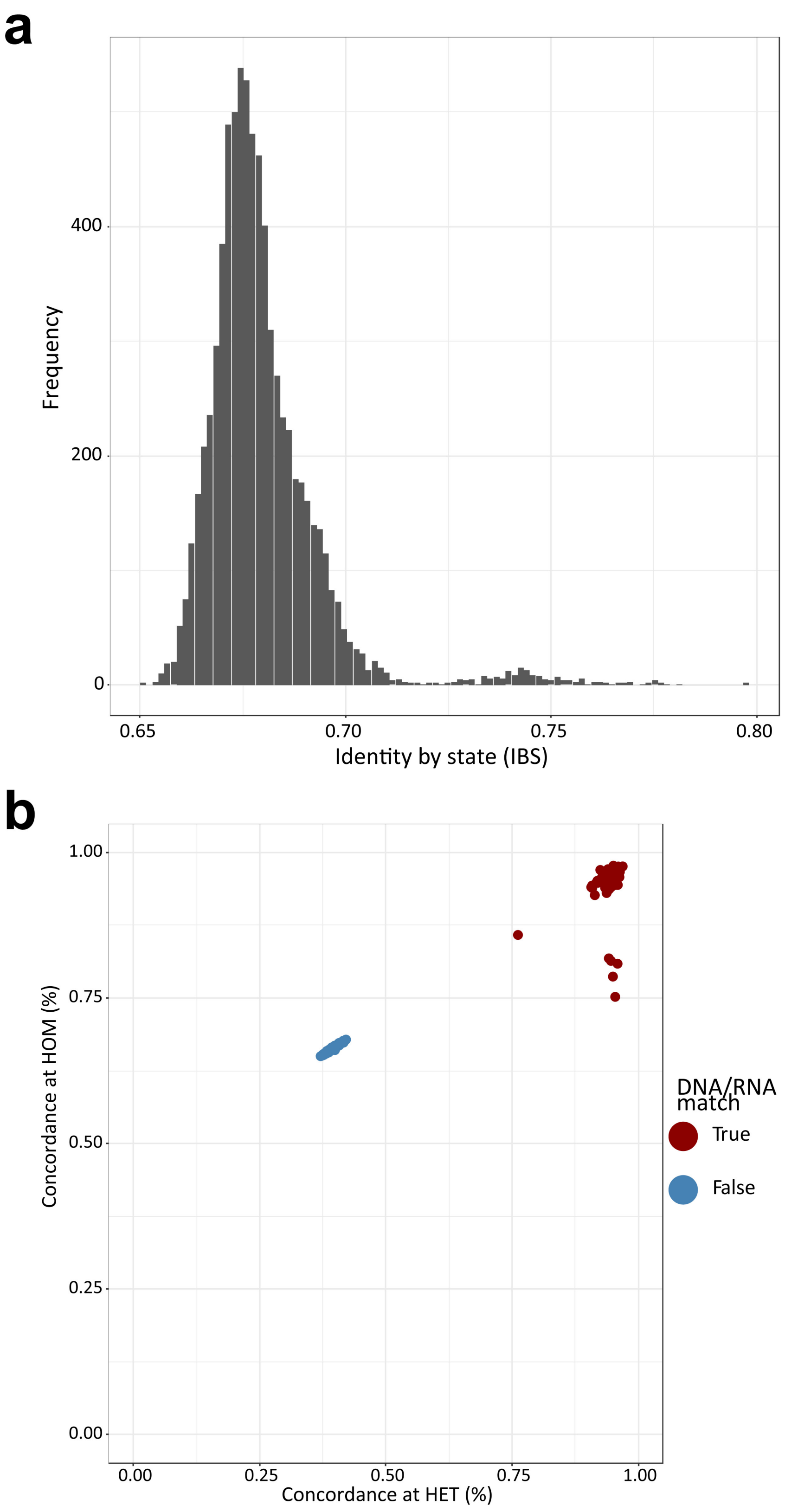

**Fig. S3: Sample duplication and DNA/RNA match assessment. a** Identity by state (IBS) values from PLINK[12] for all pairwise comparisons among the $n$ = 123 animals in the current study using the pruned SNP data prior to imputation. **B** Match BAM to VCF (MBV)[13] outputs showing allelic consistency for heterozygous ($x$-axis) and homozygous ($y$-axis) variants, between genotype and RNA-seq sequences. Red dots indicate proper identity match of RNA-sequencing with expected genotypes. As a control, we show consistency levels for non-matching samples between biological sample T064 and everything else (blue dots).

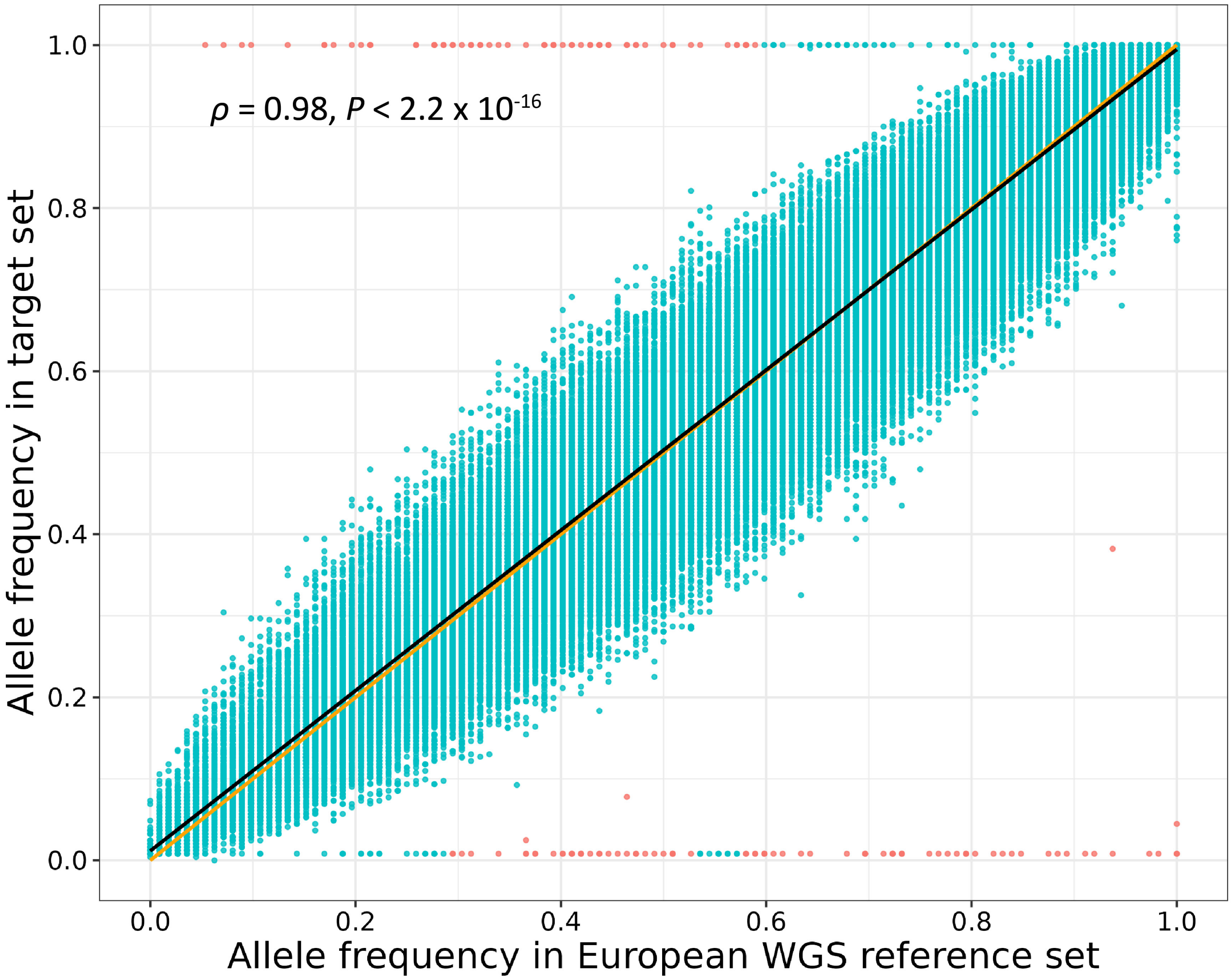

**Fig. S4: Allele frequencies between the study population and European *B. taurus* cattle from the WGS reference panel.** The observed (study-specific; *n* = 123) allele frequencies reported on the *y*-axis are plotted against allele frequencies derived from *n* = 56 European WGS animals on the *x*-axis at 236,325 matched genomic loci. Orange line denotes the line of parity. Black line denotes line of best fit. Red points indicate those removed from the study population and Global Reference Panel prior to imputation.

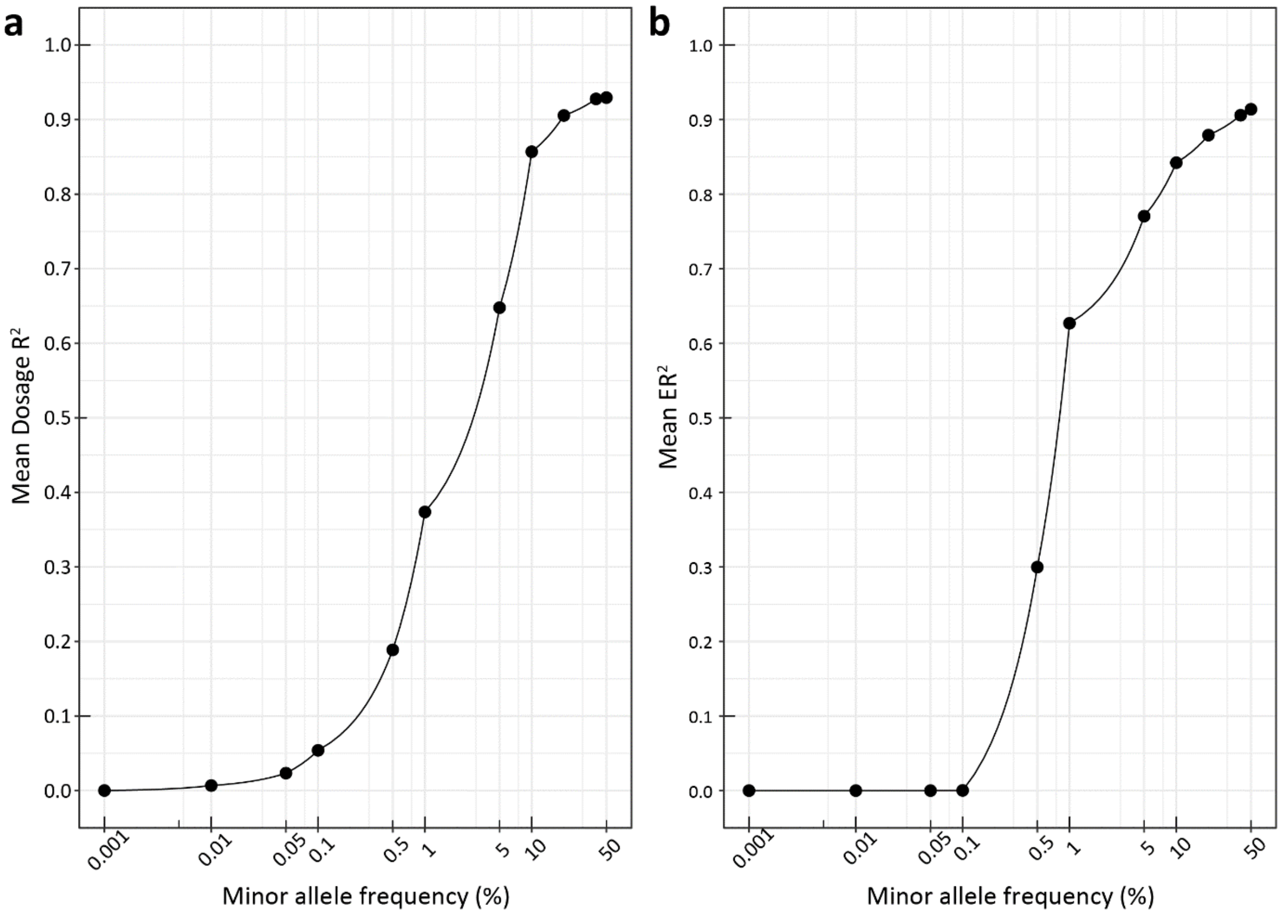

**Fig. S5: Imputation Performance. a.** Imputation accuracy (Dosage $R^2$) from Minimac4 [10] for all variants (genotyped and ungenotyped) when imputed up to WGS level using the Global Reference Panel. **b.** Imputation accuracy measured using the empirical $R^2$ ($ER^2$) from Minimac4 of genotyped variants when imputed up to WGS level using the Global Reference Panel.

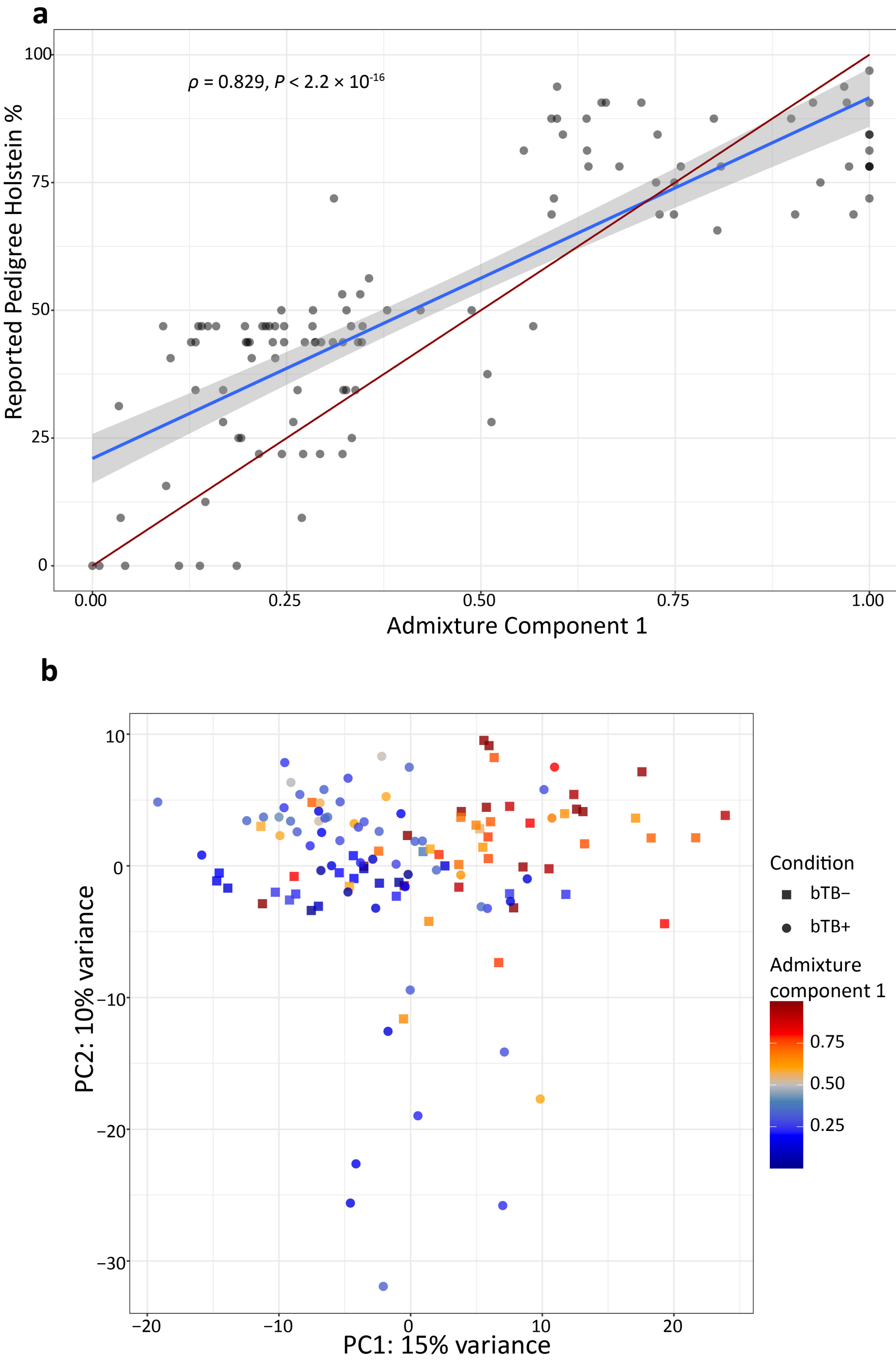

**Fig. S6: Admixture analysis and its impact on the transcriptomics principal component analysis (PCA). a** Correlation between reported pedigree Holstein % and Admixture component 1 from Admixture analysis [14] of 34,272 pruned SNPs. Red line indicates line of parity, blue line indicates line of best fit. **b** Principal component analysis (PCA) of top 1,500 most variable genes for all $n$ = 123 animals after variance stabilising transformation using DESeq2 [15]. Principal components PC1 and PC2 are plotted. Animal data points are shaped based on their experimental condition and coloured according to their corresponding Admixture component 1 value.

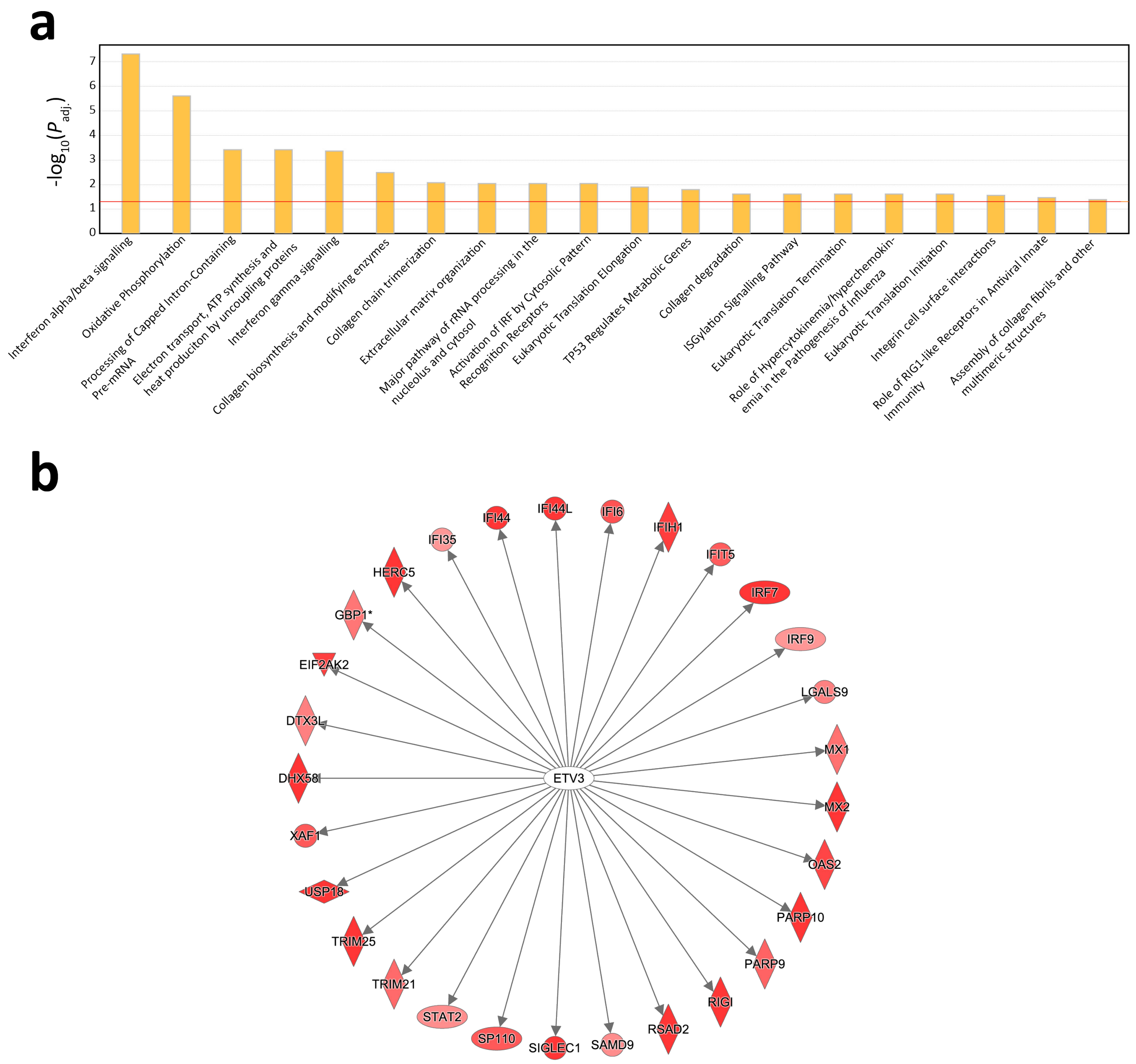

**Fig. S7: IPA® enrichment and upstream regulator results for differentially expressed genes. a** Bar plot of significantly enriched (FDR $P_{adj.} < 0.05$) Ingenuity® Pathway Analysis (IPA) pathways for highly significant (FDR $P_{adj.} < 0.01$) DE genes used as input. Bars are ordered corresponding to their -$\log_{10} P_{adj.}$ value. **b** Top upstream biological regulator (ETV3) predicted to be upregulated by IPA based on the highly significant DE genes used as input.

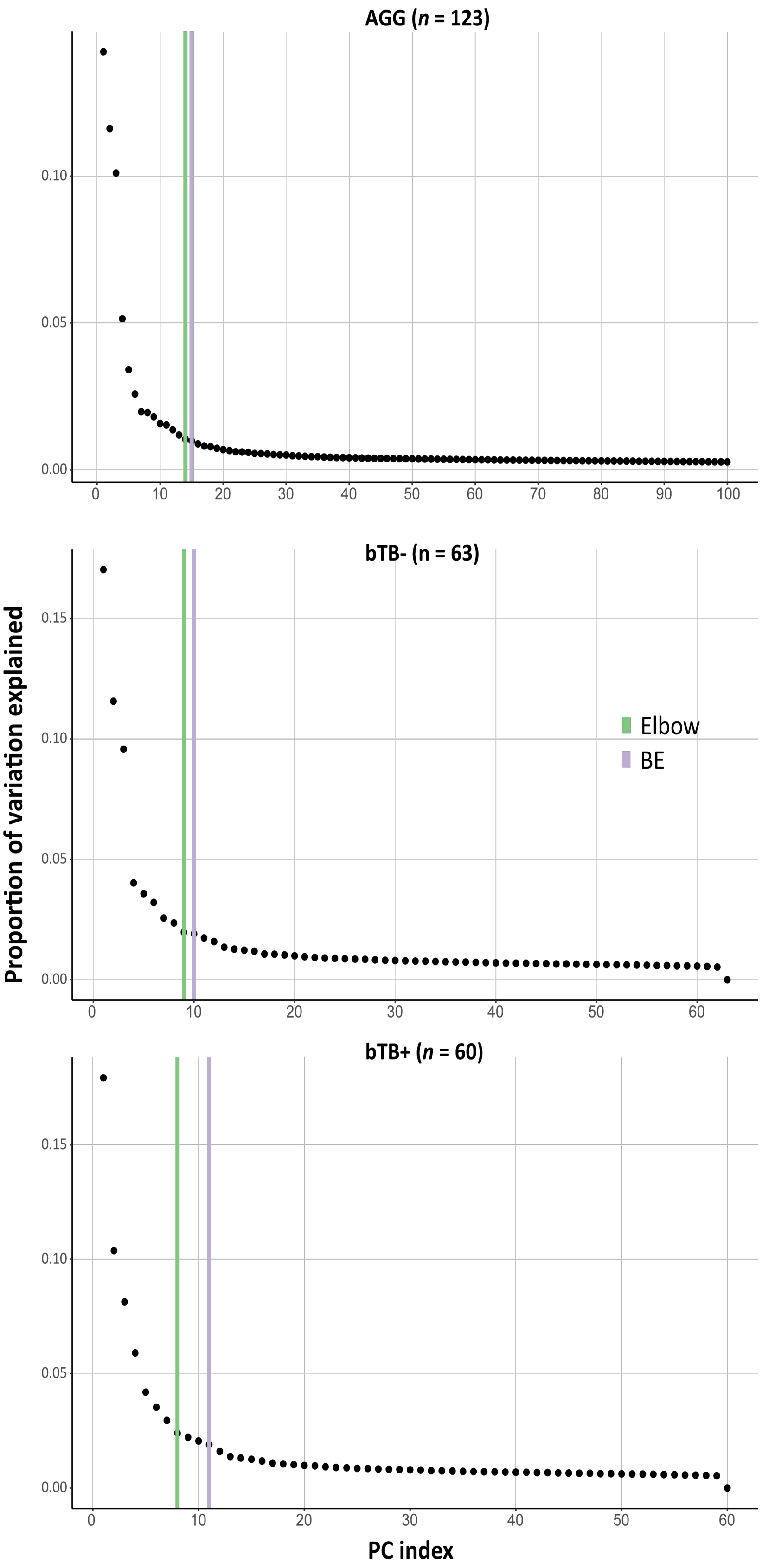

**Fig. S8: Transcriptomics principal components (PCs) used in the eQTL analysis.** Scree plot showing the proportion of variation explained by each of the transcriptomics PCs identified using the PCAForQTL R package [16] across the all animals group (AGG), the control group (bTB−), and the reactor group (btB+), respectively. Green lines correspond to the number of transcriptomic PCs inferred by the elbow method. Purple line denotes the number of PCs inferred by the Buja and Eyuboglu (BE) algorithm [17].

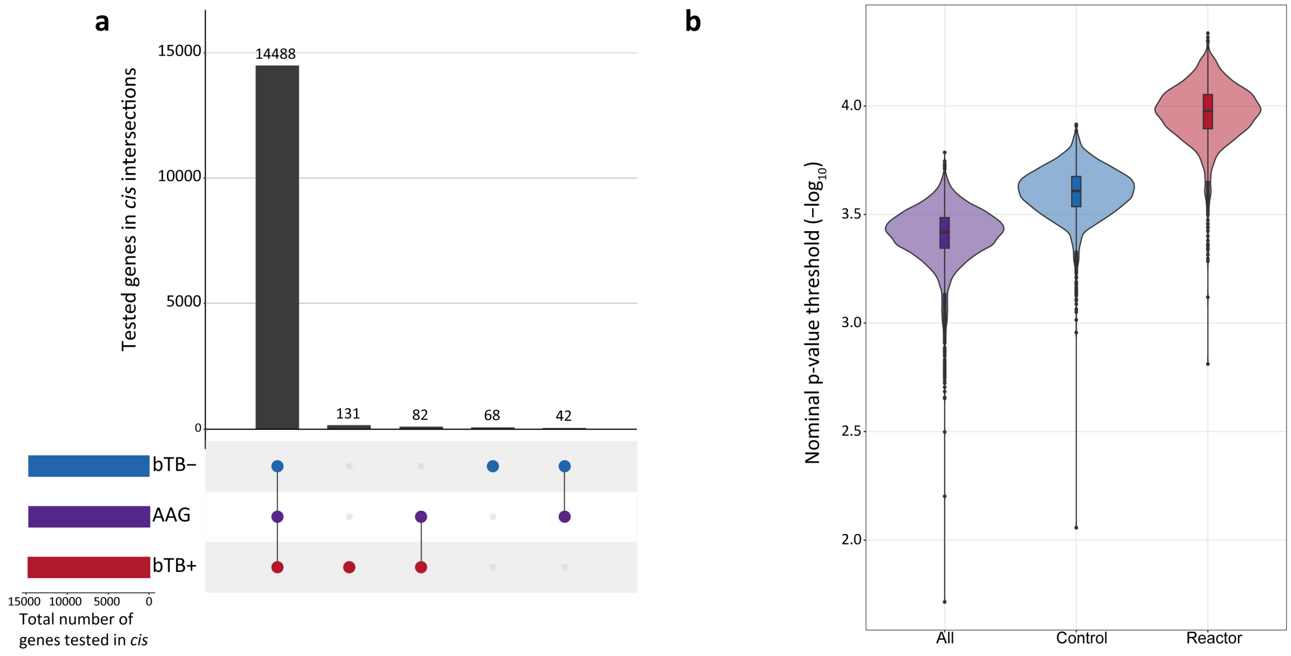

**Fig. S9: Comparison of genes tested across all three groups in the *cis*-eQTL analysis and nominal P-value thresholds within each group for the determination of *cis*-eVariants. a** Upset plot showing the number and intersection of genes tested across the bTB−, bTB+, and the AAG groups in the *cis*-eQTL analysis. **B** Violin plots showing the distribution of nominal *P*-value threshold per *cis*-eGene for bTB− (control), bTB+ (reactor), and AAG (all animal) groups, respectively.

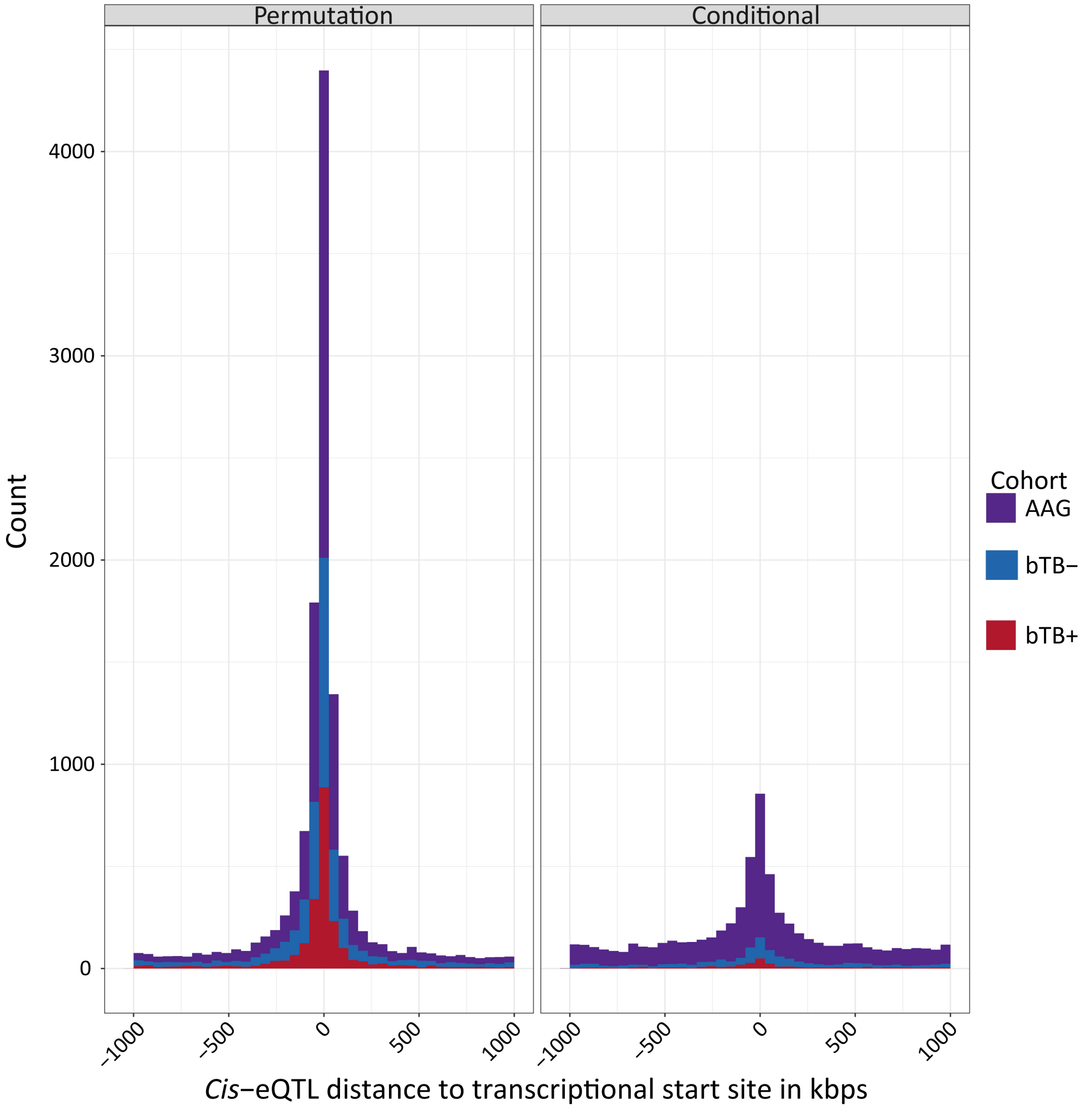

**Fig. S10: Distance to transcriptional start site of *cis*-eQTLs.** Comparison of the distances to transcriptional start site of all cis-eQTLs identified via the permutation and conditional analysis across the control (bTB-), reactor (bTB+), and combined all animals (AAG) cohorts.

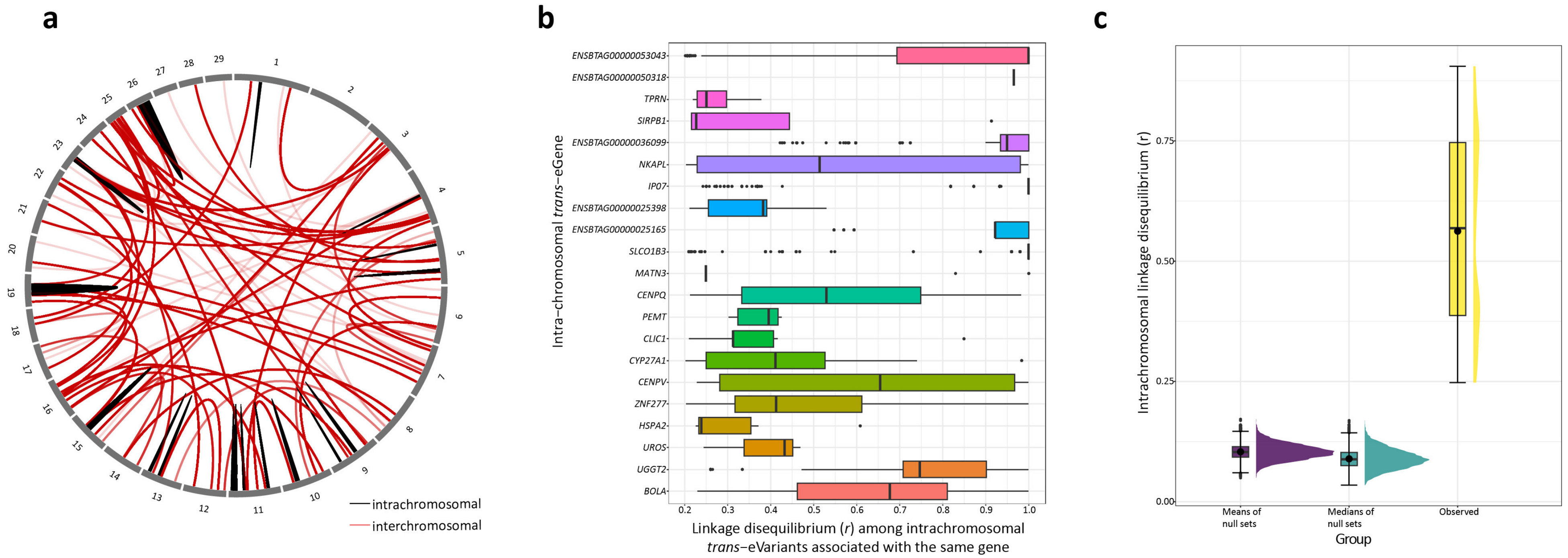

**Fig. S11: Visualisation of *trans*-eQTL analysis results and integrative intrachromosomal *trans*-eQTL analysis. a** Circos plot showing interchromosomal (red) and intrachromosomal (black) *trans*-eVariant:gene associations in the all animals group (AAG). **b** Boxplots showing the linkage disequilibrium (LD) relationship (*r*) among intrachromosomal *trans*-eVariants associated with the same gene. **c** Comparison of observed intrachromosomal LD relationship (*r*) between 24 top intrachromosomal *trans*-eVariants and *cis*-eQTLs of the same gene versus the mean and median distributions of 10,000 sets of 24 intrachromosomal null variant pairs at least 5 Mb and no more than 14 Mb away from each other. Horizontal lines inside the boxes show the medians, solid circles show the means. Box bounds show the lower quartile (Q1, the 25th percentile) and the upper quartile (Q3, the 75th percentile). Whiskers are minima (Q1 – 1.5 × IQR) and maxima (Q3 + 1.5 × IQR) where IQR is the interquartile range (Q3-Q1).

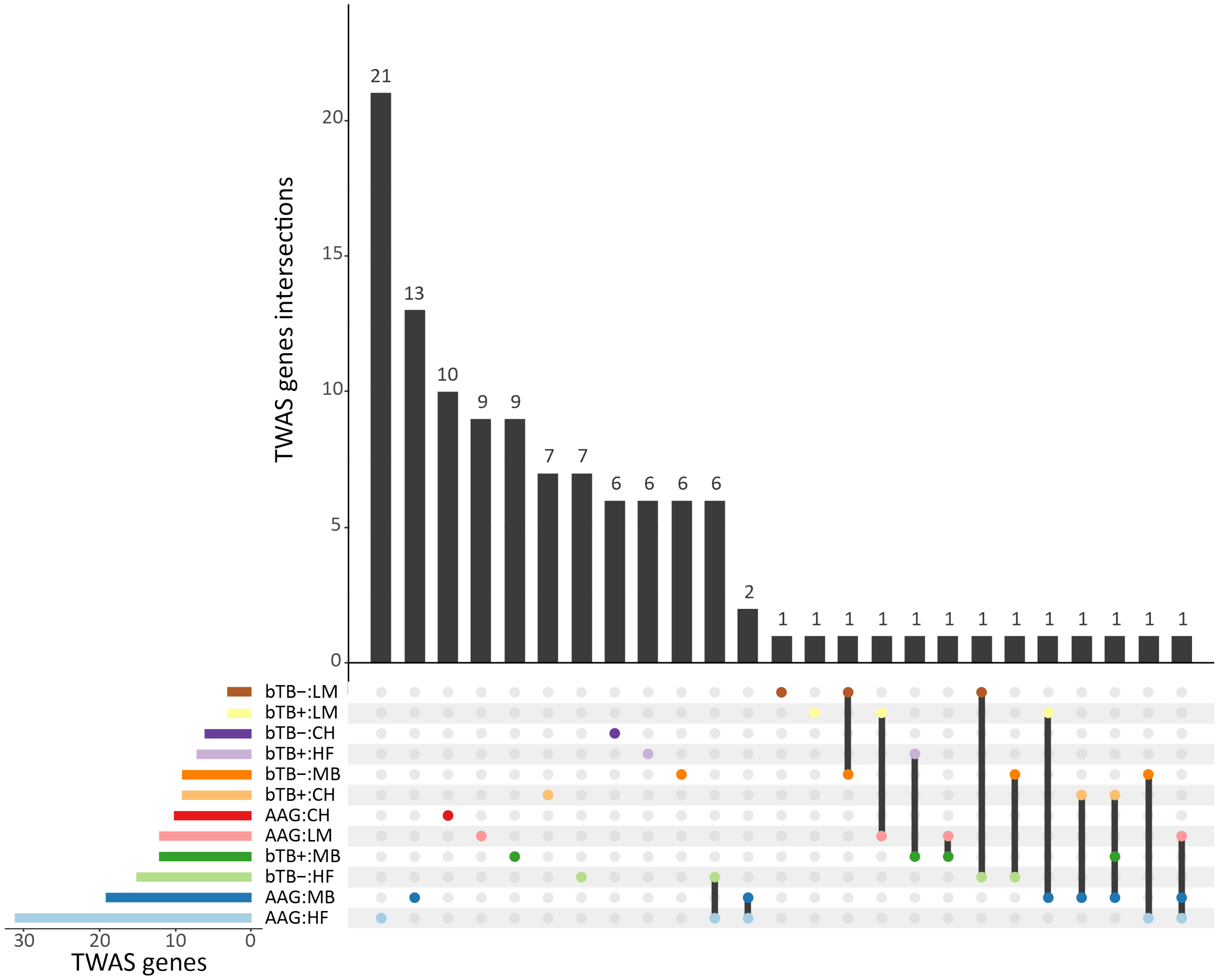

**Fig. S12: Overlap of significant TWAS genes across all 12 TWAS groups.** Upset plot showing the number and intersection of significant transcriptome wide association study (TWAS) genes (Bonferroni $P_{adj.} < 0.05$ and $P < 0.05$ post-permutation) across the bTB−, the bTB+, and AAG reference panels identified using the Charolais (CH), Limousin (LM), Holstein-Friesian (HF), and Multi-breed (MB) GWAS data sets.
